# A programmable seeker RNA guides target selection by IS*1111* and IS*110* type insertion sequences

**DOI:** 10.1101/2024.04.26.591405

**Authors:** Rezwan Siddiquee, Carol H. Pong, Ruth M. Hall, Sandro F. Ataide

**Affiliations:** School of Life and Environmental Sciences, The University of Sydney, NSW 2006, Australia

## Abstract

IS*1111* and IS*110* insertion sequence (IS) family members encode an unusual DEDD transposase type and exhibit specific target site selection. The IS*1111* group include identifiable subterminal inverted repeats (sTIR) not found in the IS*110* type [1]. IS in both families include a noncoding region (NCR) of significant length and, as each individual IS or group of closely related IS selects a different site, we had previously proposed that an NCR-derived RNA was involved in target selection [2]. Here, we found that the NCR is usually downstream of the transposase gene in IS*1111* family IS and upstream in the IS*110* type. Four IS*1111* and one IS*110* family members that target different sequences were used to demonstrate that the NCR determines a short seeker RNA (seekRNA) that co-purified with the transposase. The seekRNA was essential for transposition of the IS or a cargo flanked by IS ends from and to the preferred target. Short sequences matching both top and bottom strands of the target were identified in the seekRNA but their order in IS*1111* and IS*110* family IS was reversed. Reprogramming the seekRNA and donor flank to target a different site was demonstrated, indicating future biotechnological potential for these systems.

## INTRODUCTION

The majority of autonomous mobile elements found in bacteria that use a transposition mechanism, i.e. insertion sequences (IS) and transposons, exhibit little or no target site selectivity [3, 4]. A few examples of highly selective target site and orientation specificity that involve additional proteins have been studied [5–9], and recently a new family of IS that exhibit target site and orientation specificity using a single IS-encoded protein has been identified [10, 11]. However, the flexible target selection seen in the IS*110* and IS*1111* families has not been investigated. IS in these families encode an unusual transposase type, referred to as DEDD transposases due to the presence of an N-terminal RuvC-like catalytic domain that includes the DEDD residues [12, 13], but each member or group of closely-related members exhibit specificity for a different site where they are found in only one orientation [4, 14]. The founding IS of this type, IS*110*, was identified in the 1980s [15] and soon acquired relatives [3]. Later, IS related to IS*1111* [16] which encoded more distantly related transposases and had further distinguishing features were proposed to form a separate family [17]. However, to date how the IS in these groups identify the appropriate target and move to it has received little attention [14, 18].

A comprehensive search for IS related to IS*1111* conducted 20 years ago and detailed analysis of over 50 IS identified confirmed the proposal that there were at least two families headed by IS*1111* and IS*110* [1]. A similar comprehensive analysis of the IS*110* group is not available. IS*1111* family members recovered from available sequences could be distinguished simply from IS*110* family members via the presence of sub-terminal inverted repeats (here designated sTIR), usually 11-13 bp in length with a perfect or near perfect match. The terminal extensions that are part of the IS but found beyond the inverted repeats (IR) were asymmetric and later work has shown that their correct lengths are 7 bp on the left and 3 bp on the right [2, 19] or occasionally 6 bp and 3 bp. In addition, the transposases aligned well with those of IS*110* and its relatives in the N-terminal catalytic (DEDD, RuvC) domain but less well at the C-terminus [1]. The presence of a non-coding region (NCR) of significant length downstream of the transposase gene (*tnp*) in all cases was noted [1], and later, it was proposed that this NCR was involved in selection of the appropriate target site [2]. However, although this feature is not found in any of the IS families encoding DDE transposases, the NCR is generally not mentioned in later descriptions of these IS [4, 14, 18]. Over time, both families have grown but, despite the clear differences, they are currently grouped together under “IS*110* family” in the ISFinder database [20] (www-IS.biotoul.fr) and discussed together in reviews of IS sequences (e.g. [18]). Indeed, as the features of and relationships within the IS*110* type have not been examined in detail, there may be further families embedded in this group.

Experimental characterisation of IS in these two families is limited. However, members of both the IS*1111* and IS*110* families have been shown to form a circular intermediate in which the IS ends are directly abutted [1, 2, 13, 21–25]. This creates a promoter that in a few cases has been shown to be active [22, 23]. This promoter is presumed to drive transcription of the *tnp* gene and would also transcribe the NCR. In a few cases, transposition to the appropriate target has been demonstrated for both IS*1111* family IS [1, 22, 23] and IS*110* family IS [13, 21, 26]. Though a duplication of bases in the target sequence has been claimed, in other cases no duplication was found. In fact, a duplication of target sequences does not appear to be characteristic of IS in either family and this discrepancy likely results from incorrect identification of the ends of the IS which should include one copy of the duplication. Further information on what has previously been reported about these families can be found in the online resource TnPedia in TnCentral [14].

Here, we have first confirmed the separation of the IS*1111* and IS*110* families showing that the longer NCR is upstream of the *tnp* gene in IS*110* family members and examining the phylogeny of a curated set of predicted transposase sequences. We then examined the role of the NCR in IS movement and in selecting the correct target. We used several IS*1111* family members, ISEc11 [22], which moves to a short linear target, ISKpn4 and ISPst6 which both target one end of certain *attC* sites of integron-associated gene cassettes [2, 23], and ISPa11 which targets the REP sequences in *Pseudomonas aeruginosa* chromosomes [1]. We also examined one IS*110* relative, ISEc21, for which the target sequence was known but its location was not and this was determined here. We have shown that seeker RNA (seekRNA) determined by the NCR found downstream of the *tnp* transposase gene in the IS*1111* ISs and upstream of *tnp* in the IS*110* IS, is essential for movement and co-purifies with the transposase. The seekRNA was found in a long form and a more abundant short form that both include a short region complementary to each strand of the target. In the seekRNAs of IS*1111* family IS, the top strand complement precedes the bottom strand match but, for the IS*110* family IS, the order was reversed. Hence, the mechanics of target selection may be different in the two families. Instances of natural reprogramming were identified and we reprogrammed the system for a representative of both IS*1111* and IS*110* families to insert a mini IS carrying a cargo gene into chosen sites.

## RESULTS

### IS*1111* and IS*110* family features

Here, we noticed that the longer NCR is upstream of the *tnp* gene in IS*110* and some of its relatives rather than downstream as in the IS*1111* type (Fig. 1a). Hence, over 350 IS listed in ISFinder (www-IS.biotoul.fr), currently under the designation IS*110* but grouped as IS*110*, IS*1111* or unknown type, were examined to locate the longer NCR. 186 had the NCR downstream of *tnp*, 123 upstream, 26 had two longer NCRs and 17 had none. Almost all correctly classified IS*1111* family IS, i.e. those with sTIR of sufficient length and a transposase related across the full length to that of IS*1111*, had the longer NCR downstream of the *tnp* gene. However, a few had an upstream NCR or a large NCR both up and downstream. In the remaining IS, the NCR was generally upstream, and sTIR were not present or were short and imperfect. However, some of these had a 3’-NCR. Hence, the NCR location appears to be a characteristic of the two families, albeit with some exceptions. This finding supports the conclusion that the IS*1111* family is indeed distinct from the IS*110* family and that the mechanisms used by these two families may differ significantly, particularly with regard to end recognition.

**Fig. 1.**
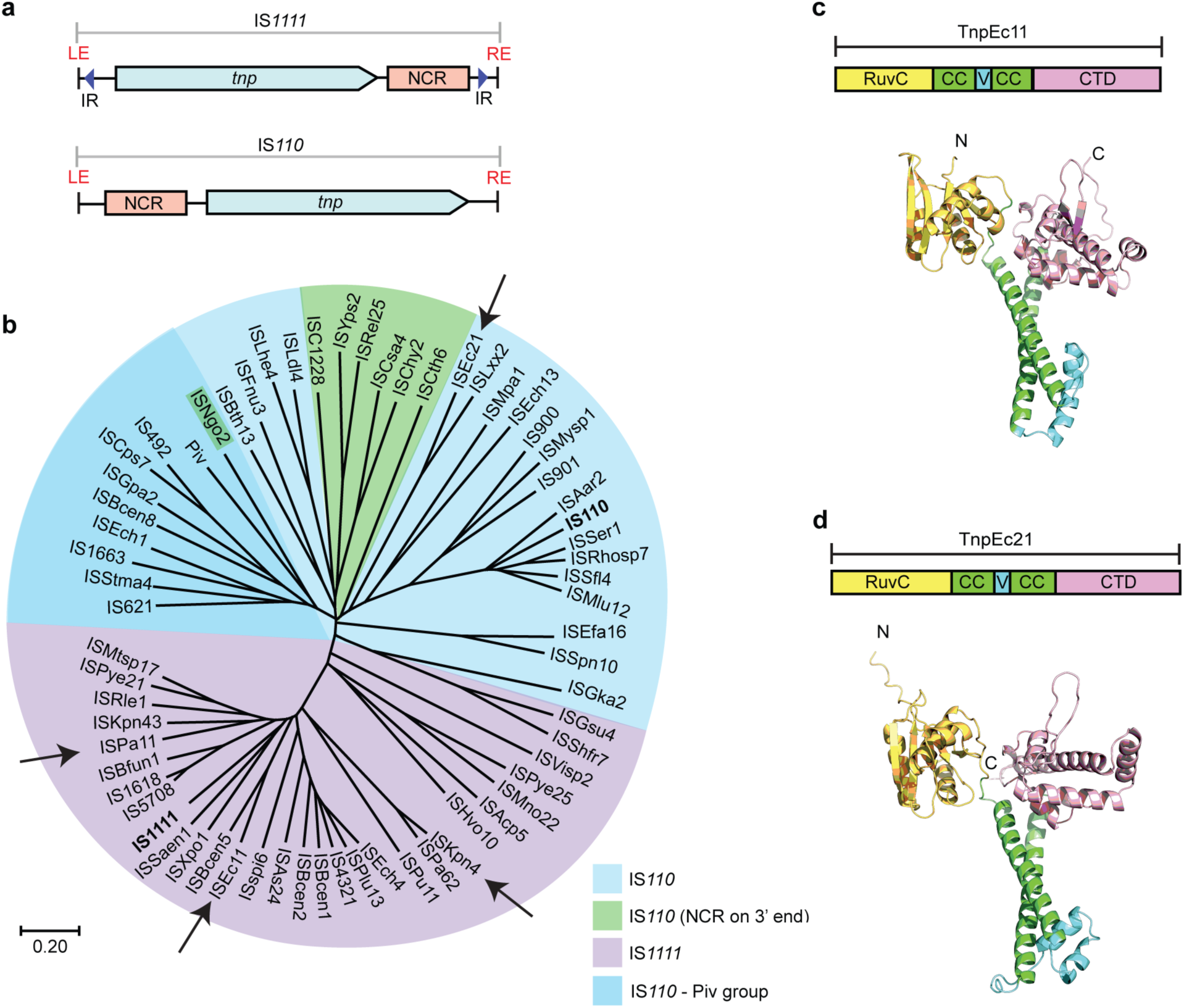
IS*1111* and IS*110* family features. **a.** Organization of IS*1111* and IS*110* family members showing locations of the longer non-coding region (NCR, pink box) relative to the transposase gene, *tnp* (blue arrow). Subterminal inverted repeat (sTIR) in IS*1111* ISs are indicated by arrows. **b.** Simplified phylogenetic tree of a few transposase sequences from IS*1111* and IS*110* family members and Piv, rooted by midpoint. The scale bar represents a 20% divergence in protein sequence. Founding family members IS*110* and IS1111 are indicated in bold. Arrows point to ISs studied in this work. Colour code key is indicated. AlphaFold 2 model prediction of domain structures of **c.** TnpEc11 and **d.** TnpEc21 as examples of IS*1111* and IS*110* family transposases, respectively. The RuvC domain (yellow), a coiled-coil domain (green), a variable region (cyan) and an uncharacterized C-terminal domain (CTD) (pfam: PF02371) (pink) are highlighted.

Previous studies had found that the N-terminal catalytic domain of the transposase in both IS*110* and IS*1111* relatives is related but the C-terminal domain aligns poorly [1, 13, 27] and our alignments (Extended Data Fig. 1) confirmed this. While in the logo for the IS*1111* family transposases a clear pattern of consensus residues is apparent in both the N- and C terminal regions, only an SG (part of a G--P---SG motif) is highly conserved in the C-terminus for the IS*110* family transposases. For the IS*1111* transposases, this location is represented by SG or TG in a GL-P----S(t)G motif. Conservation of the SG residues in a G..P….S(t)G motif was noted previously [13, 27], and these two aa were shown to be required for inversion of the invertible DNA segment of *Moraxella* catalysed by Piv [27], which is a relative of the IS*1111* and IS*110* family transposases [28].

A phylogeny of the transposases constructed from the set of IS listed in ISFinder, curated to include only a single representative for each group of transposases with >70% pairwise aa identity (Extended Data Fig. 2), revealed that the transposases of IS*1111* family members, classified based on the presence of sTIR of appropriate length, are clearly separated from the remaining transposases from IS*110* members. Two exceptions (marked in blue) are found in a deep branching group of transposases from IS classified as IS*110* family. A simplified phylogenetic tree (Fig. 1b) highlights some further exceptions. In the tightly clustered group of IS*1111* relatives all but one had a downstream NCR. In ISMtsp17 the NCR was upstream, in contrast to its closest relative ISPye21 which had a standard downstream long NCR. The more diverged IS, ISHvo10 and ISAcp5, have long NCRs both up and downstream. For the IS in the IS*110* family, the NCR was found mainly upstream of *tnp* but it is downstream in some e.g. the group highlighted in green in Fig. 1b. Transposases represented in the dark blue group in Fig. 1b that includes IS*492* [21] and IS*621* [13] are most closely related to the Piv inversion protein [12, 28]. The phylogeny (Extended Data Fig. 2) also indicates that there are clear subfamilies within each family and there may be further families particularly in the current IS*110* group granted the large range of lengths (315-450 aa) found in transposases encoded by members of this family compared to the more compact spread of lengths in the IS*1111* family transposases (330-350 aa). For example, transposases in the branch of the IS*110* family that includes the IS*492*, IS*621* and the Piv invertase (Extended Data Fig. 2) are generally much shorter at 315-330 aa than the transposases of members of the subgroup that includes IS*110* which range from 395-455 aa. The transposases of IS in the cluster that includes ISEc21 used in this study are of intermediate size (345-385 aa).

However, there are clear similarities in the folded structures of the transposase proteins. Using AlphaFold2 [29], the structures of several transposases derived from each family were modelled, and transposase TnpEc11 from ISEc11 (339 aa), as a representative for the IS*1111* family, and TnpEc21 from ISEc21 (358 aa), as a representative of the IS*110* family, are shown in Fig. 1c and Fig. 1d, respectively. The transposases all consist of an N-terminal DEDD (RuvC fold) catalytic domain [28] and a C-terminal domain with no well characterised homologues (includes Pfam PF02371) separated by a coiled coil with a variable region between the α-helices. More detailed representations of these transposases showing the configuration of the DEDD residues and the location of the SG motif at the tip of the finger in the C-terminal fold are in Extended Data Fig. 3. The boundaries of the various domains are marked on the conservation analysis for TnpEc11 and TnpEc21 derived using the Consurf server (consurf.tau.ac.il) (Extended Data Fig. 4). The variable region, which is composed of small α-helices and loops, is longer in the ISEc21 transposase than in that of ISEc11. AlphaFold2 also predicts the formation of a dimer via interactions between the coiled coils placing the variable region of one monomer next to the RuvC domain of the second monomer (Extended Data Fig. 3). The AlphaFold2 modelling also predicts a tetramer as shown only for TnpEc11 in Extended Data Fig.3.

Further transposases derived from the other IS*110* subfamilies were modelled and the variation in transposase lengths seen in the subgroups of the IS*110* family transposases was largely accounted for by variation in the length of the variable region. This region consisted of a simple loop in Tnp492 and Tnp621 from the Piv group, and in Tnp110 there is a larger region of helices and loops than in TnpEc21. However, the precise role of the variable region is not currently known.

### The downstream NCR in ISEc11 is essential

ISEc11 is an IS*1111* family member that was reported to recognises a short target (GTGAAAATACTG) and has been shown to form a circular intermediate in which an active promoter is generated at the junction and to transpose to its preferred target [22]. Here, ISEc11 (cloned in pUC19 together with approximately 100 bp of flanking sequence) produced a circular intermediate that was readily detected using PCR (Fig. 2a). However, the uninterrupted target sequence that would be regenerated if the flanking sequences are re-joined (as in a site-specific tyrosine or serine recombinase reaction) was not detected under the same conditions, indicating that the IS is not simply excised. However, when the 3’-NCR was deleted (bp 1170-1374 of 1443 bp removed) leaving the transposase gene and right end intact, the circular form was not produced consistent with an essential role for this region. Moreover, when the 270 bp fragment that includes the NCR (bp 1127-1397 of ISEc11 preceded by a T7 promoter and followed by an HDV ribozyme) was supplied in a separate plasmid, the circular form was again produced (data not shown). Likewise, transposition of ISEc11 to its preferred target in a separate plasmid required the presence of the NCR *in cis* or *in trans* (Fig 2b). In the product, the IS was inserted into the target at the appropriate position and in the correct orientation, and as expected for IS*1111* family members no additional bases were generated (Fig. 2c). Note that, previously, a 4 bp target site duplication was claimed for this IS [22] but here one copy of the 4 bp was assigned to within the IS at the left end to form the required 7 bp extension at that end. Replacement of target bases flanking the donor on either side prevented both circle formation and transposition (Fig. 2d), demonstrating that intact target-derived sequence on both sides of the donor is needed for formation of the circular intermediate.

**Fig. 2.**
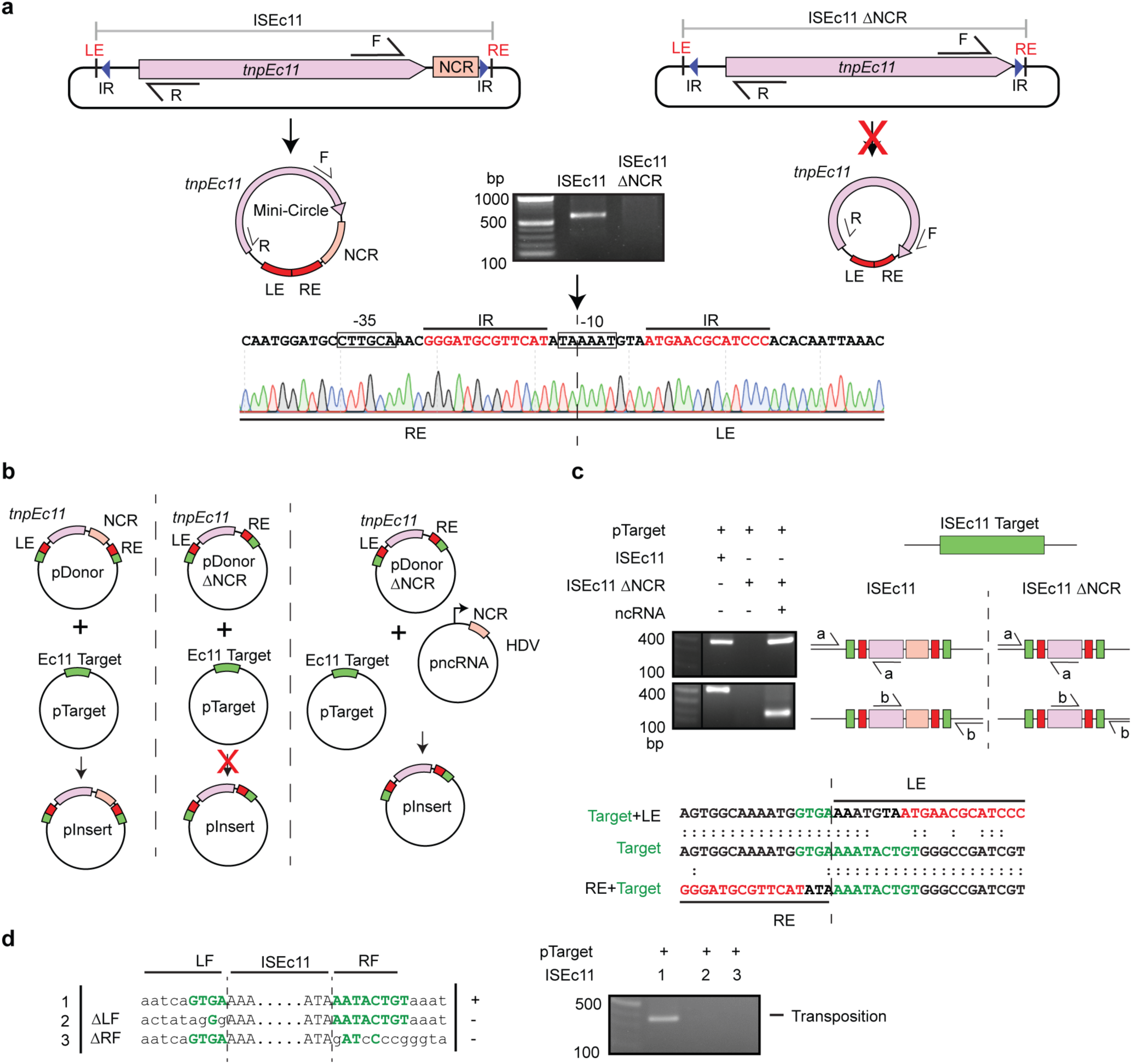
Features of ISEc11, an IS*1111* family member, required for transposition. **a.** The plasmids containing ISEc11 or ISEc11ΔNCR, lacking the non-coding region (bases 1170-1374 of 1433), flanked by the ISEc11 target are shown above predicted circular intermediates with an arrow indicating whether the reaction occurred. Outward facing primers R and F used to detect circular intermediates are indicated on the plasmids. Sanger sequencing of the R-F PCR products formed for intact ISEc11 is shown below with the promoter motifs formed by joining the left and right ends (LE and RE) boxed and the IR in red type. The junction is indicated by a vertical arrow and dashed line. **b.** The *in vivo* transposition assays demonstrating the requirement for the NCR. Plasmids present are indicated above the vertical arrow. The predicted product is below the arrow. pTarget contains the ISEc11 target (green), pDonor contains ISEc11 flanked by the target, pDonor contains ISEc11 lacking the NCR and pncRNA contains the NCR. The pInsert plasmid shows the potential transposition products with ISEc11 or ISEc11ΔNCR inserted within the target site. **c.** PCR detection of *in vivo* transposition. Primer pairs (a and b) used to detect transposition are indicated on a schematic of potential produces. Sequences of the PCR products compared to the target are shown below with the IR in red and target in green. confirms the correct target site insertion point. **d.** Altered flanking target sequences tested. Consensus target bases are green. PCR products formed using primers R and F are shown to the right.

### IS*1111* family transposases co-purify with NCR-derived RNA

The ISEc11 transposase, TnpEc11, was expressed (using a HisMBP and Strep TagII fusion), either with or without the full length ISEc11 present in the same cells to supply the NCR, and then purified. The yield of TnpEc11 was poor in the absence of the NCR region but improved when the complete ISEc11 (preceded by a T7 promoter) was present to supply the NCR (Fig. 3a). The TnpEc11 expressed in the presence of the NCR purified with a nucleic acid and distinct bands were clearly present. The nucleic acid was digested by RNase but not by DNase (Fig. 3b). RNA extracted from the affinity purified RNP complex (Figs. 3c) was sequenced, and the reads were aligned with the ISEc11 sequence where they mapped to the 3’-NCR (Fig. 3d). Hence, RNA transcribed from the essential NCR was specifically associated with the transposase.

**Fig. 3.**
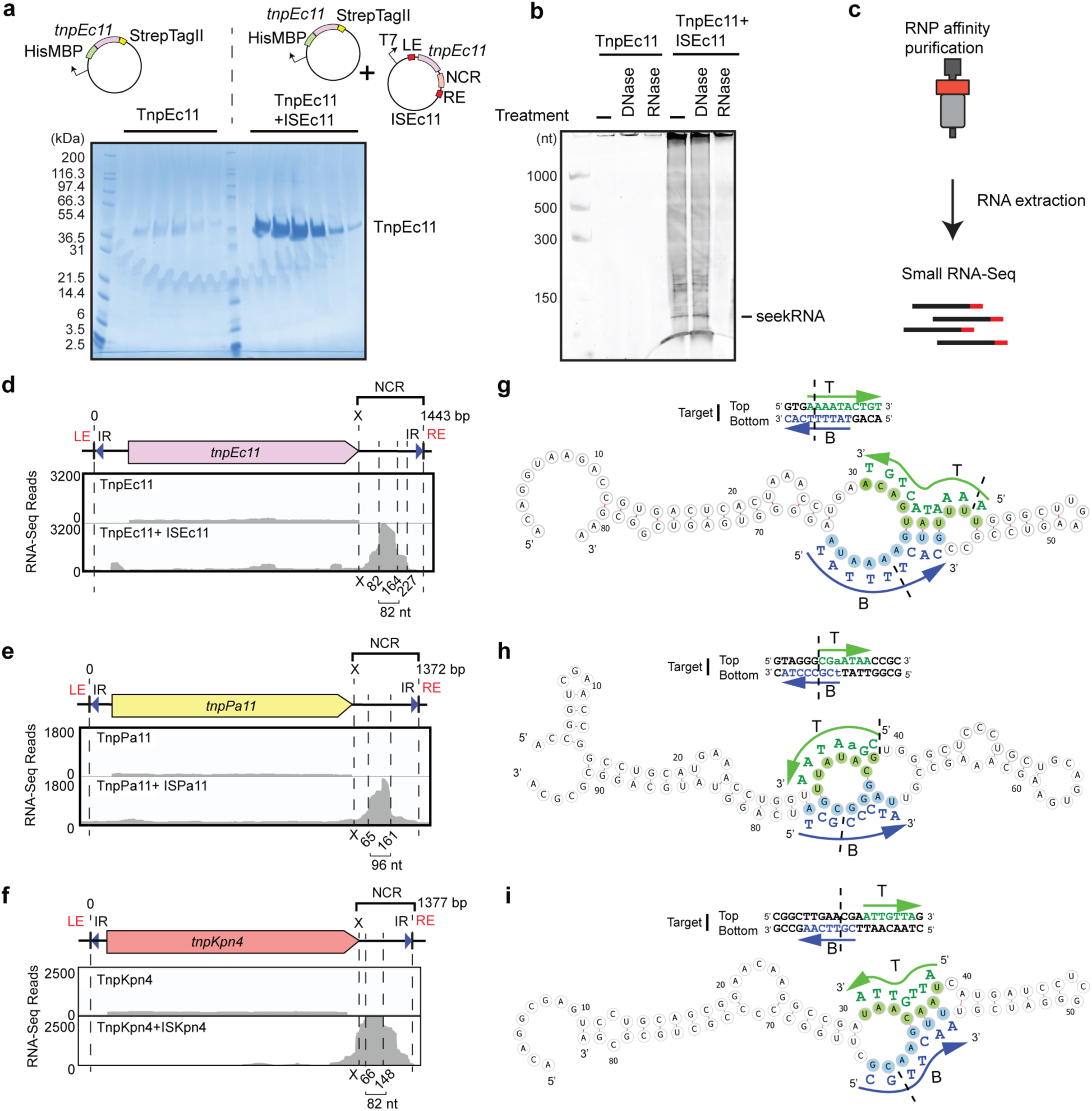
Characterisation of seeker RNA bound to of IS*1111* family members. **a.** SDS-polyacrylamide gel electrophoresis (PAGE) of TnpEc11 purified alone or in the presence of ISEc11, showing TnpEc11 yields were greatly enhanced by the presence of the NCR. **b.** SYBR GOLD-stained denaturing PAGE of TnpEc11 purified alone or in the presence of ISEc11 after digestion with DNase or RNase. **c.** Schematic of the RNP and RNA purification and RNA sequencing steps. **d.** Small RNA-seq reads mapped to ISEc11 with the sizes of the longer seekRNA and the peak, shorter seekRNA below, together with their location relative to the *tnp* termination codon (x). **e.-f.** As in d. for ISPa11 (e) and ISKpn4 (f). ISPa11 (e) and ISKpn4 (f) sequence. **g.** Folded structure prediction of the ISEc11short, peak seekRNA with the target logo above. Sequences that base pair with the top (T) strand of the target are highlighted in green and with bottom (B) strand of the DNA target are blue. Arrows indicate the 5’-3’of the target DNA strand and dashed lines indicate the insertion point of the IS. **h.-i.** As in g. applied to ISPa11 (h) and ISKpn4 (i).

To confirm that this property is found more widely in IS*1111* family members, two further IS that target different sites were examined. ISKpn4 and ISPa11 have previously been shown to form a circular intermediate and are found at a specific location in a potentially folded structure, an *attC* site of integron-associated gene cassettes for ISKpn4 [2] and a *P. aeruginosas* REP for ISPa11 [1]. As for ISEc11, TnpKpn4 and TnpPa11 both purified with an associated RNA (Extended Data Fig. 5) and this RNA mapped to the region downstream of the *tnp* gene (Figs. 3e and f).

### A seekRNA finds the target

To locate the target-determining region or regions, we searched in the sequence of the predominant 82 nt RNA associated with TnpEc11 for short sequences that could base pair with the forward and/or the reverse strand of the target sequence. Two short segments, one complementary to the forward or top strand (T in Fig. 3g) and one complementary to the reverse or bottom strand (B in Fig. 3g), were found, and their location in the folded structure of the RNA, predicted using MXfold2 [30], is shown in Fig. 3g. These matches overlap in the target (Fig. 3g). As these matches can explain the ability to select a specific target, the NCR-derived RNA was named a seeker RNA or seekRNA. The folded structure for the longer seekRNA is in Extended Data Fig. 6. The sequences of the seekRNA corresponding to the predominant peak for ISPa11 (96 nt) and ISKpn4 (82 nt) were also examined and matches located in the predicted folded structure (Fig. 3h and 3i). Again, two short segments complementary to the forward and the reverse strand of the target sequence were found in the sequence of the predominant band and were in the same order as in ISEc11 seekRNA, namely top strand match located 5’-to the bottom strand match. Predicted folds for the corresponding long seekRNAs are in Extended Data Figs. 6 and 7.

ISPst6 inserts into the same position in a target the same or very similar to the ISKpn4 target [23] and TnpPst6 is 86% identical (92% similar) to TnpKpn4. ISPst6 also produced a seekRNA that purified with the transposase (Extended Data Fig. 5) and the ISPst6-derived 86 nt seekRNA contained the same matches as the ISKpn4 seekRNA (Extended Data Fig. 7). We also examined the NCR of ISPa25 that also targets this site but encodes a significantly diverged transposase (TnpKpn4 and TnpPa25 are 46% identical; 60% similar) [2]. The region in the DNA sequences where they most closely match includes the region corresponding to the short seekRNA of ISKpn4 (Extended Data Fig. 7). The predicted seekRNA was found to include the same stretches of sequence matching the target and assume a very similar fold (Extended Data Fig. 7). Hence, the seekRNA does not appear to be confined rigorously to interaction only with a specific group of very closely related transposases (>70% identity).

### The seekRNA is programmable

ISEc11 and ISXne4 represent an example of natural re-programming of the seekRNA. When the predicted short seekRNA of ISXne4, a relatively close relative of ISEc11 (TnpEc11 and TnpXne4 are 72% identical; 88% similar), was compared to the seekRNA of ISEc11 (Extended Data Fig. 8), a high level of identity was observed but the target-matching bases in the ISEc11 seekRNA were not found (Fig. 4a). Examination of the location of the few known copies of ISXne4 revealed that it was surrounded by a different sequence and that this target matched the differing regions (Fig. 4a). This confirms that the two target-matches have been correctly identified and demonstrates re-programming.

**Fig. 4.**
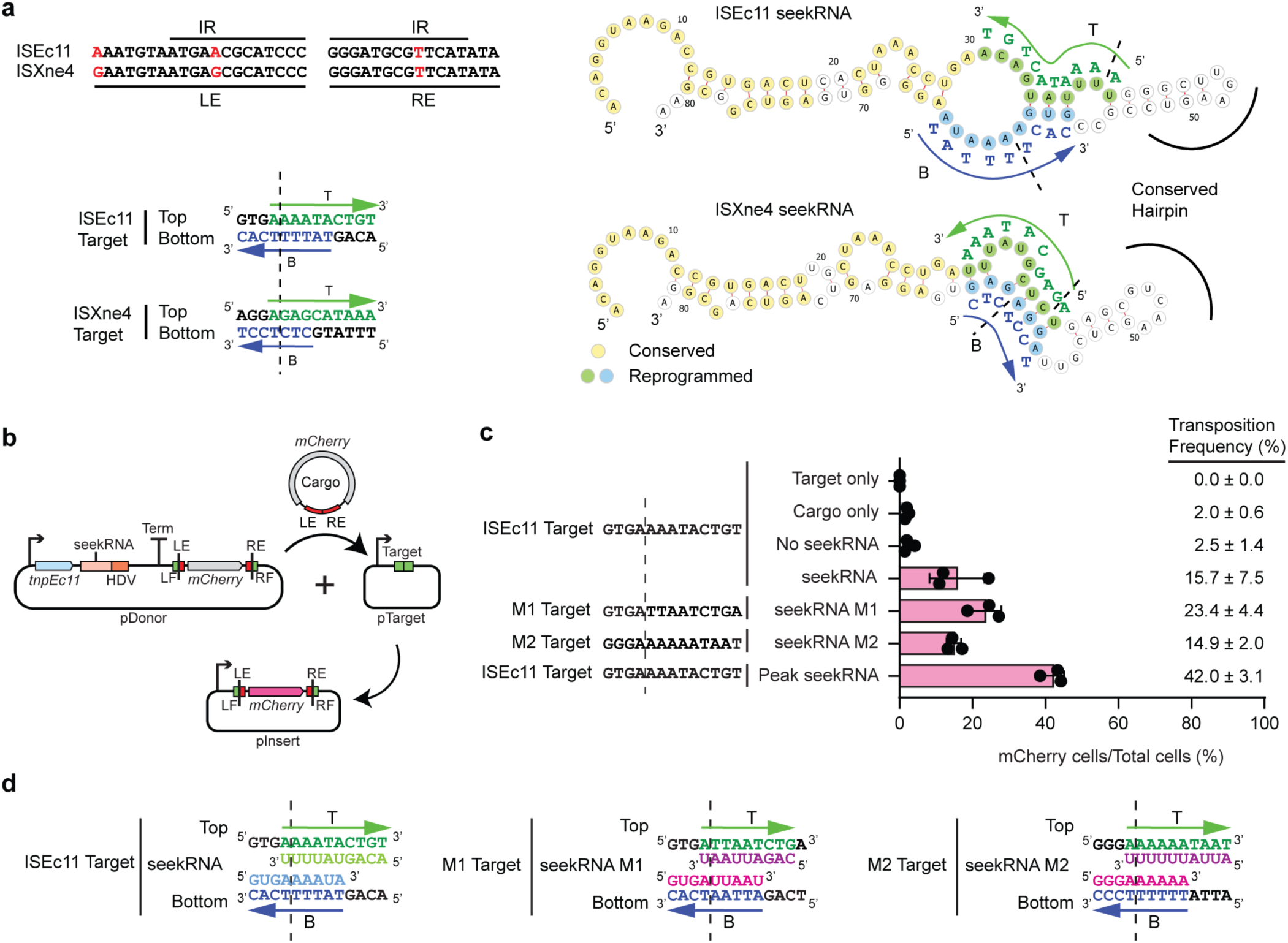
Natural and experimental reprogramming to recognize a new target site. **a.** Features of ISEc11 and ISXne4 indicating natural reprogramming. On left above, an alignment of the outermost left and right IS ends (LE and RE) with differences in red and sTIR overlined. Below, the target sites for ISEc11 and ISXne4. A dashed vertical line indicates the IS insertion point. Arrows labelled T and B indicate the bases with complementary sequences in the seekRNA of ISEc11 and ISXne4. On the right, folded structure predictions for ISEc11 and ISXne4 seekRNA with nucleotides conserved in both in yellow, nucleotides complementary to their target site in green or blue and a conserved hairpin indicated. The bases in the target are shown alongside in green and blue with an arrow indicating the 5’-3’ direction. **b.** Schematic of the mCherry reporter assay. The pDonor plasmid contains a T7 promoter preceding *tnpEc11* and the natural or re-programmed seekRNA followed by a HDV ribozyme with a strong T7 terminator at the 3’end. The mini IS contains mCherry without promoter surrounded by the LE and left flanking (LF) sequence and the RE and right flanking sequence (RF). pTarget includes a target (LF and RF abutted) preceded by a T7 promoter (bent arrow). In each assay, the LF and RF in the donor and target plasmids are identical and appropriate to the matches in the seekRNA. **c.** Transposition efficiency of the mCherry mini IS by TnpEc11 with the natural (ISEc11) or reprogrammed seekRNA (M1 and M2) or the peak ISEc11 seekRNA. The target sequences used are shown with vertical dashed line marking the insertion point. Black dots indicate triplicate experimental determinations with standard deviation shown as bar. Transposition frequencies are on the right. **d.** Sequences of the ISEc11 target and modified targets (M1 and M2) with the base-pairing sequences present in the seekRNA below the top strand sequence and above the bottom strand sequence. Arrows point to the 3’of the DNA target site in both strands T (top) and B (bottom). Dashed line indicates the insertion point of the target site.

To re-programme the IS to move to a different target, it was necessary to change both the sequences corresponding to the target that flank the donor IS and the target matching sequences in the seekRNA. For ISEc11, two new targets were tested using movement of a mini IS containing the mCherry gene without an upstream promoter bounded by the IS ends (LE 50 bp, RE 46 bp) and flanks containing the target (Fig. 4b). The TnpEc11 and the appropriate long (154 nt) seekRNA were supplied in the donor plasmid. Movement to targets (Fig. 4c and Extended Data Fig. 9) that were preceded by a T7 promoter enabled expression of the mCherry, measured by FACS sorting (Extended Data Fig. 10). Using the wildtype target and seekRNA, transposition occurred at a frequency of about 15% (Fig. 4d). When the portion of the target that flanks the IS on the right was altered and the corresponding changes were made in the seekRNA (see Extended Data Fig. 9 for details), transposition of the mCherry to the new M1 target occurred at about 23% frequency. In the second case, the target on both sides of the donor mCherry mini IS and corresponding positions in the seekRNA were altered, and again the transposition frequency to the new M2 target was 15%. Hence, the long seekRNA could be programmed to move the IS to a different location. Using the same assay, the short 82 nt seekRNA (peak in Fig. 3d) was also tested and the mCherry mini IS moved even more efficiently (42% transposition; Fig. 4d, bottom line) indicating that this length is sufficient to support IS movement.

### Moving cargo

To demonstrate that a cargo gene or genes can be moved by these IS, the *catA1* chloramphenicol resistance gene with its upstream promoter and ribosome binding site (adding 745 bp) was introduced at the start (after bp 50) and at the end (after bp 1397) of ISEc11 (Fig. 5a). In both cases, movement was detected. The central portion of ISEc11 was also replaced by the *catA1* gene with an upstream promoter leaving the ends intact. When the transposase and NCR were supplied in trans, again movement was detected (Fig. 5b) as seen for mCherry, indicating that the system can be harnessed to mobilise different cargos.

**Fig. 5.**
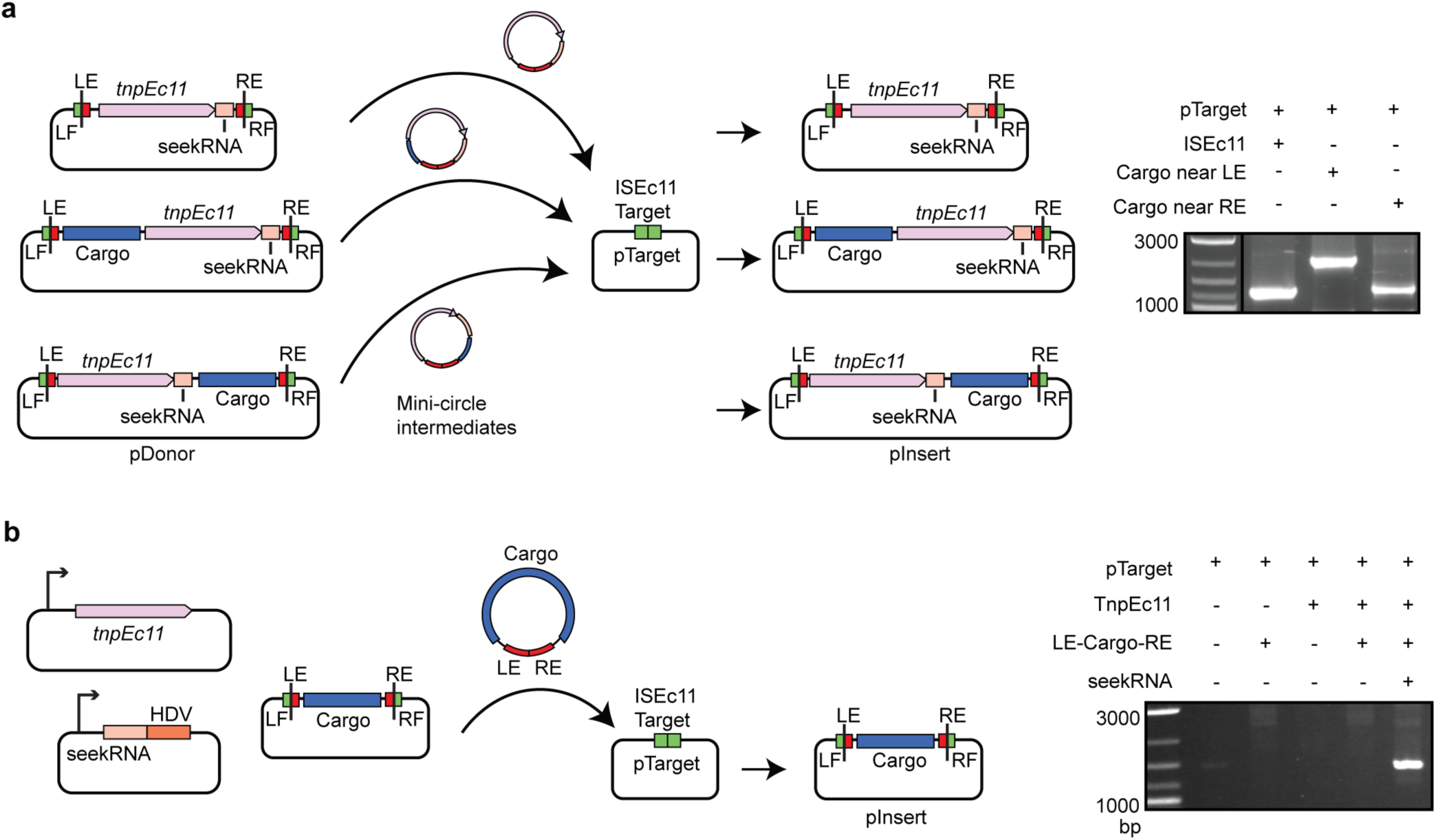
ISEc11 can carry cargo. **a.** Schematic of IS transposition assay. The pDonor plasmid includes ISEc11 or ISEc11 with the *catA1* gene (blue box) labelled cargo inserted upstream of the *tnp* after 50 bp from the start of the IS or downstream of the *tnp* 46 bp from the end of IS and pTarget included the ISEc11 target (green box). Movement to pTarget was detected by PCR (right panel) using primers in pTarget and in *tnp*. The mini-circle intermediate is also shown. **b.** Schematic of mini IS transposition assay. For minicircle formation three compatible plasmids are present. The mini IS contains the *cat* cargo flanked by 84 bp of from the left IS end (LE) and 71 bp from the right end (RE) and is flanked by target sequences (4 for LF and 7 bp for RF). The *tnpEc11* and the long seekRNA followed by a HDV ribozyme, each under the control of the T7 promoter (bent arrow), are in separate plasmids. Movement of the mini IS into a fourth plasmid, pTarget containing the ISEc11 target, was detected by PCR (right panel) using a primer in *catA1* and a primer in pTarget and PCR products were sequenced.

### Similarities and differences for ISEc21 from the IS*110* family

ISEc21 is an IS*110* family member with an upstream NCR (Fig. 6a). The target listed in ISFinder was confirmed computationally and here determined to be a linear target after it was traced to a conserved region within certain IS*3* family members (e.g., ISCfr6, ISEc92, ISEc93), namely the codons for and surrounding the second D of the DDE in the catalytic domain of the transposase (Extended Data Fig. 11). As for ISEc11, ISEc21 formed a circular intermediate in which the ends were abutted and a promoter generated (Fig. 6a). When the upstream NCR was deleted (bp 20-150 removed), circle formation no longer occurred (Fig. 6a). The NCR was also needed *in cis* or *in trans* (bp 8-179 in ISEc21 preceded by a T7 promoter and followed by an HDV ribozyme) for transposition (Figs. 6b and 6c) and movement into the target supplied at the correct position and in the correct orientation without generating a duplication of bases at the target site was detected (Fig. 6d). When the number of target bases matching the consensus was reduced to only 3 on the right or to 5/6 on the left and 5 on the right of the donor IS, both minicircle formation and transposition still occurred (Fig. 6e). However, removal of further bases on each side abolished movement, again indicating the importance of the presence of the two parts of the target sequence surrounding the donor IS for recognition of the IS ends and to bring them together in the minicircle.

**Fig. 6.**
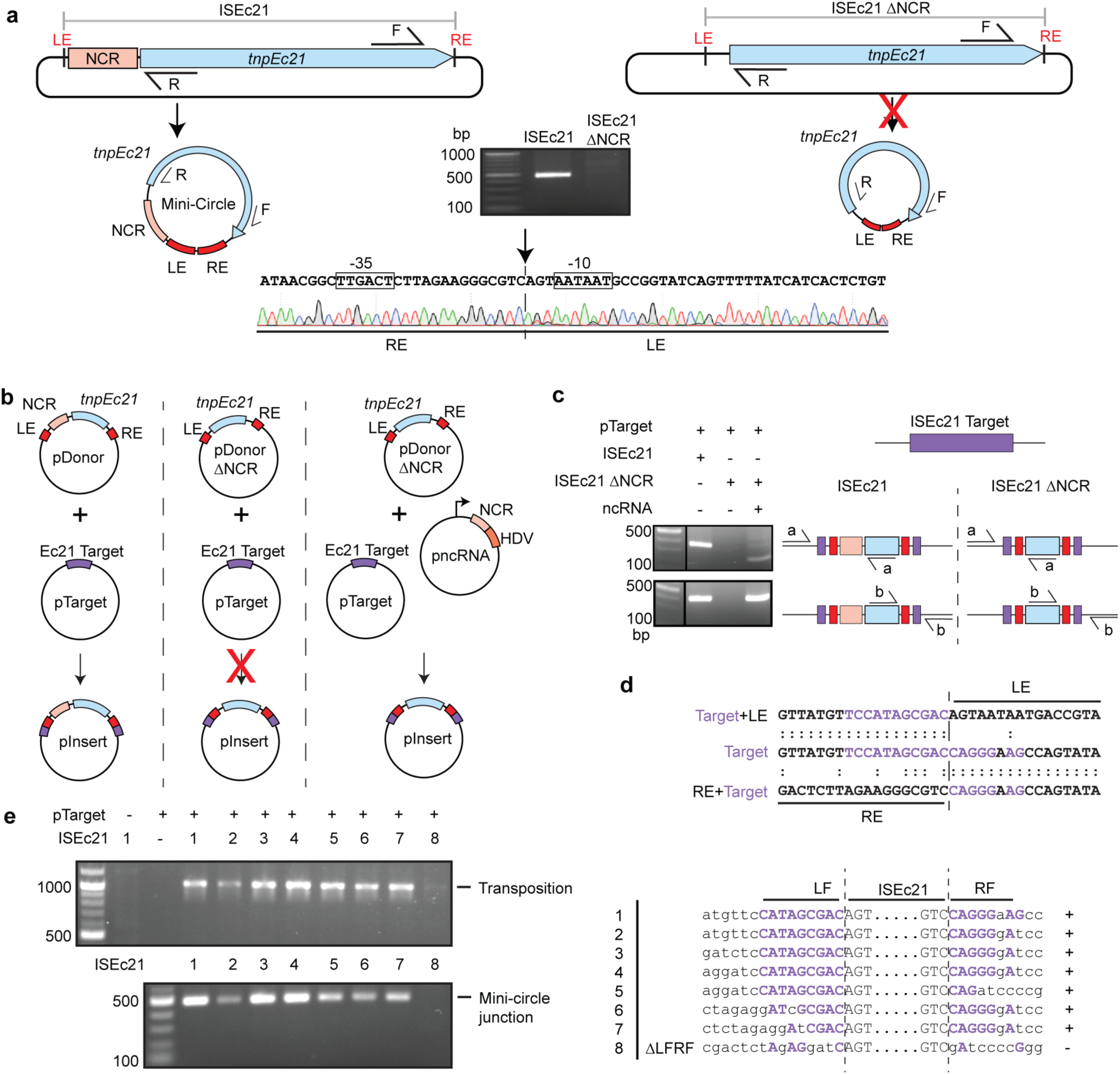
Features of ISEc21, an IS*110* family member, required for transposition. **a.** The upstream ncRNA is required for transposition. The plasmids containing ISEc21 and ISEc21ΔNCR, lacking the non-coding region (bases 20-150), are flanked by the ISEc21 target. They are shown above predicted circular intermediates with an arrow marked to indicate if the reaction occurred. Outward facing primers R and F in the *tnp* gene used to detect circular intermediates are indicated on the plasmids. R-F PCR products and Sanger sequencing of the PCR products formed for intact ISEc11 is shown below with the promoter motifs formed by joining the left and right ends (LE and RE) boxed and the junction indicated by a vertical arrow and dashed line. **b.** Transposition assay configurations. The plasmids shown in a. combined with a compatible plasmid containing the target (pTarget) or the ISEc21ΔNCR plasmid with pTarget and a third plasmid carrying the NCR followed by and NDV ribozyme and preceded by a T7 promoter (bent arrow) are shown above an arrow marked to indicate if the product (pInsert) below the arrow was formed. **c.** Detection of *in vivo* transposition. Primer sets a and b complementary to the target plasmid and the *tnp* gene at each end of the IS are shown schematically on the right with the PCR products shown on the left. **d.** Sequence of PCR products containing the LE/RF and RE/RF junctions compared to the target The insertion point is indicated by a vertical dashed line. **e.** Requirement for flanking target sequences. Configurations of the sequences in the LF and RF are shown on the right. Conserved bases in the target are in purple capitals and both adjacent bases and bases altered in the target are black lower case. PCRs to detect transposition and circle formation are shown on the left.

The TnpEc21 transposase purified with RNA (Extended Data Fig.5) that was recovered and sequenced. The sequences mapped to the upstream NCR (Fig. 7a, Extended Data Fig. 12). Again, a shorter seekRNA was more abundant than the long seekRNA (see Extended Data Fig. 12 for the fold of the long seekRNA). Bases matching both strands of the target were found in the 74 nt short seekRNA (Fig. 7a). However, in contrast to the situation in the seekRNAs from all of the IS*1111* family members tested here, the top strand match (T in Fig. 7a) is located 3’- to the bottom strand match (B in Fig. 7a).

**Fig. 7.**
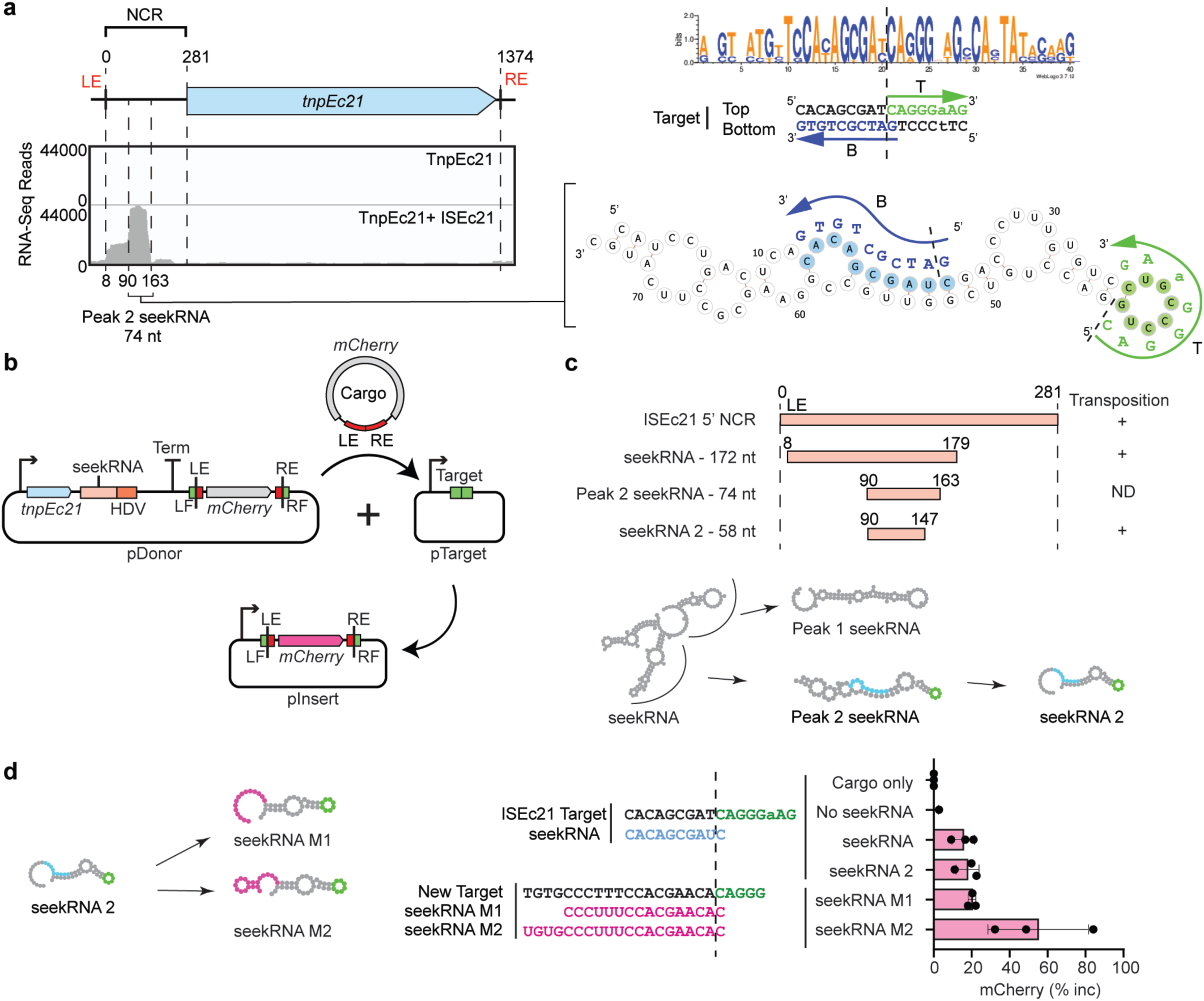
Characterisation and experimental reprogramming of ISEc21. **a.** The ISEc21 seekRNA. Small RNA-seq results from RNA purified from the TnpEc21 RNP complex aligned with the ISEc21 sequence shown schematically above. Positions of RNAs are shown together with the size of the peak seekRNA. On the right, predicted folds for the peak (small) seekRNA with sequences complementary to the target in blue and green. Complements are shown in blue and green above or below; T and B indicate top and bottom strand, respectively, and the arrow indicates 5’-3’. The target consensus is at the top with the top and bottom strands of the target below with bases complementary to the seekRNA blue and green. The dashed line indicates the insertion point. **b.** Schematic of mCherry mini IS reporter assay. pDonor contains *tnpEc21* (blue arrow) and the ISEc21 seekRNA (pink) followed by a HDV ribozyme (orange) and an mCherry mini IS. Both *tnp* and the seekRNA are expressed from a T7 promoter (bent arrow) and followed by a strong T7 terminator. The mCherry gene lacks an upstream promoter and is surrounded by the left flanking sequence containing target (LF) and LE and the RE and right flanking sequence containing target (RF). pTarget contains the target preceded by a T7 promoter. The predicted minicircle and transposition product are also shown. **c.** Activity of NCR-derived RNAs. Lengths and positions of NCR-derived RNAs tested with the outcome for transposition (+ or -) detected by PCR and confirmed by sequencing on the right. Predicted folds for the seekRNA forms are shown below. **d.** Reprogramming. New sequences are in the target-complementary regions of the shortened seekRNA, in the target and in the LF and RF of the mini IS. Transposition was detected by PCR and sequencing of the pInsert boundaries and detected using mCherry fluorescence.

Using the assay shown in Fig. 7b, a 58 nt RNA that is 16 nt shorter than the peak 74 bp ISEc21 seekRNA was found to be sufficient to support transposition. Movement of a promoterless mCherry in a mini IS, surrounded by the IS ends (LE 29 bp, RE 16 bp) with 15 bp flanks carrying the ISEc21 target site, moved to its usual target site (Fig. 7c). The system was also re-programmed. One of the target-matched regions in the shortened 58 nt ISEc21 seekRNA 2 form was changed to detect a different sequence and mCherry, was surrounded by the outer IS ends now flanked by the new target sequence (Fig, 7d Extended Data Fig. 13). Transposition occurred when the target site surrounding the donor was the same as the target offered and seekRNA included appropriate nucleotides to detect that target (Fig. 7d).

A case of natural re-programming was also found in an ISEc21 relative. The transposase of ISMch6 is 72% identical (85% similar) to TnpEc21 but the IS is found in a modified target. Comparison of the DNAs of the NCR revealed three altered bases in the target-defining region for the bottom strand (Extended Data Fig. 12) and the similarities between the predicted folded short seekRNAs for ISEc21 and ISMch6 are shown in Extended Data Fig. 12. Notably, although only a single copy of ISMch6 was found in sequences available in GenBank, the differences were found in the sequence surrounding ISMch6.

### Many IS, many different targets

The most unusual feature of the IS*1111* and IS*110* families is that the target recognised by each IS or group of IS is not the same. A computational pipeline to identify a consensus target site by searching the sequences available in the GenBank nucleotide database was developed, and applied to over 300 IS sequences found in ISFinder under the IS*110* family (https://github.com/AtaideLab/Targets). The strategy used identified all copies of each IS and then joined the flanking sequences (200 bp at each end). Redundant IS copies that were in the same broader position were removed so that only unique locations were considered (Fig. 8a). Thereafter, a consensus logo was developed for 20 bp on each side of the IS. There was enormous variation the number of copies found, ranging from one unique insertion event for 47 IS or only two for 110 IS to over 7000 unique events for IS*1663* from *Bordatella pertussis* (Fig. 8b and c). Overall, the accuracy of these logos is limited by the number of unique insertions and by the accuracy of end positions in entries in ISFinder. However, the logos obtained from the pipeline were the same as those reported previously [1] for the IS*4321* target (here with nearly 900 insertions) and ISPa11 target (here with over 1000) and for several other cases. For these, the junction between the ends and the target had been found or confirmed via comparisons with uninterrupted targets [1] and finding the same targets via the pipeline confirms the accuracy of the approach. A few logos for IS from each family are shown in Fig. 8d.

**Fig. 8.**
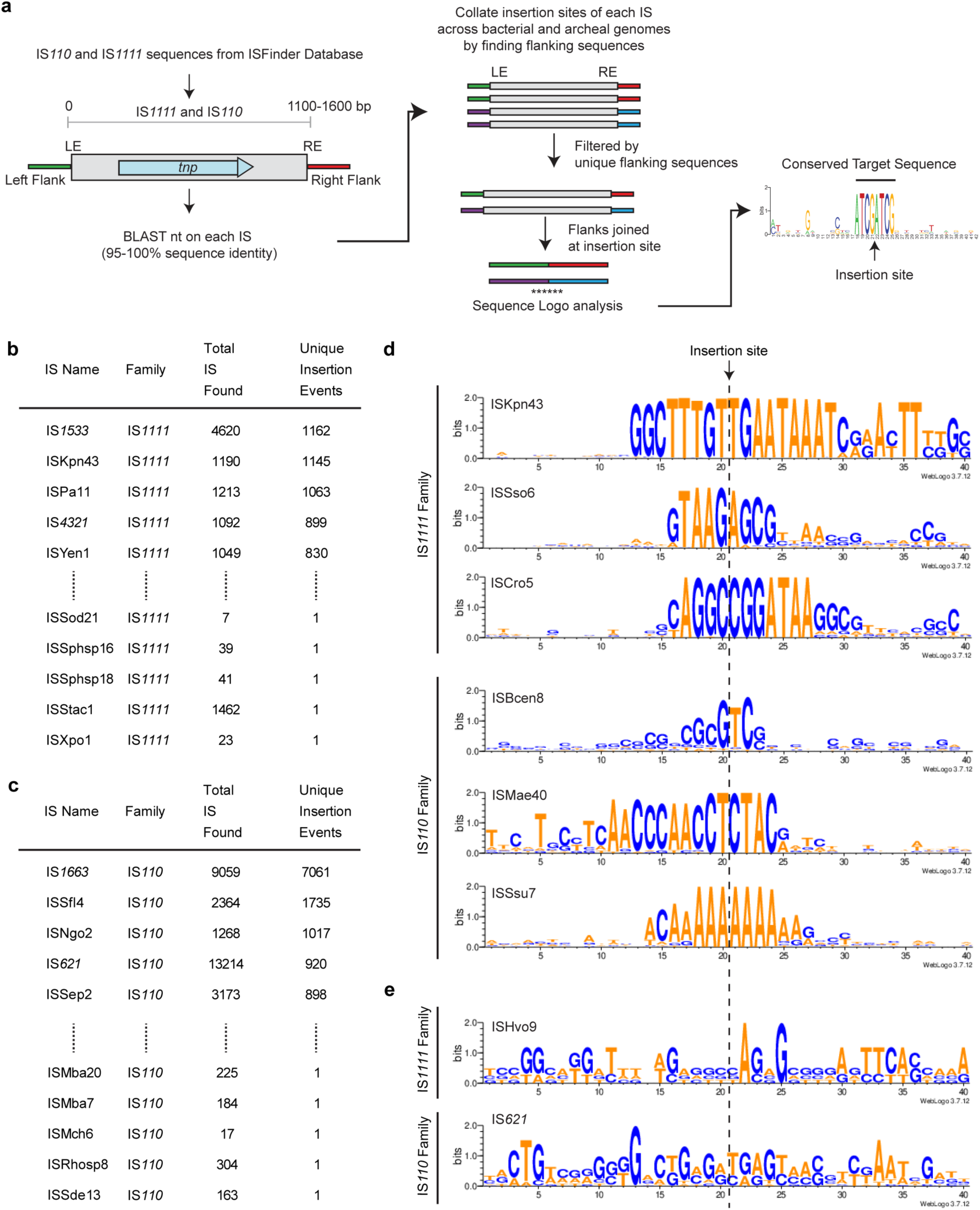
Identification of the target site of individual IS*110* and IS*1111* family members. **a.** Schematic of the pipeline and scripts used to determine the target sites of the IS IS*110* and IS*1111* family members as listed in ISFinder. The script and documentation along with a demo sample are available on https://github.com/AtaideLab/Targets/ **b.** Top and bottom 5 ISs from IS*1111* family with respective number of unique insertion events. **c.** Prevalence of IS. Top and bottom 5 ISs most abundant IS from IS*110* and IS1111 families with respective number of copies and unique insertion events. **d.** WebLogos representing the conservation in the target sites of members of IS*110* and IS*1111* families showing varied length and sequence. **e.** WebLogos showing poor conservation. An example of target site logos for one IS*1111* and one IS*110* family member showing poor conservation.

From the logos, it appears that the ends listed in ISFinder may be incorrect and need adjustment in a number of cases. In these cases, the correct target cannot be deduced computationally. For example, some early entries record adjacent duplications one of which should be in the IS and this leads to a duplication in the target logo, as can be seen in the IS*492* logo. In other cases, a conserved target was only seen on one side, indicating that conserved bases on the other side may have been included within the IS boundaries. A few early entries lack the terminal extensions found in IS*1111* family IS and the extension bases appear in the predicted target for IS*1533*, which is the most abundant IS*1111* family member (Fig. 8b). Resolution of these issues will require identification of the uninterrupted target(s), which could not be simply achieved in the pipeline developed here or a sequence of the circle junction which must be obtained experimentally.

Although in some cases a consensus was not generated because there were insufficient different locations, in others there were many locations but no clear logo (Fig 8e). In the case of Hvo9 (Fig. 8e), the target may have been included with the IS. Another example is IS*621* with 920 unique locations (Fig. 8b) but no clear consensus for the surrounds (Fig. 8e). In this case, it was possible that there are multiple versions of the IS (>95% identical) that target different sites. However, the same logo was found when the stringency in searches was raised to 100%. Clearly, manual curation will be needed to address this surprising finding.

## DISCUSSION

Here, we have shown for the first time that IS in the IS*1111* and IS*110* families that are abundant in bacteria and archaea use an RNA that we have named the seekRNA, to detect their preferred target and take up the correct position and orientation in their specified target. The seekRNA was essential and purified with the transposase as a RNP complex. It provides a flexible target site selection mechanism and natural re-programming was detected among the relatives of the IS examined here. However, the specifics of how target recognition occurs may be different in the two families as the regions that match the two strands of the target are in reverse order in the IS*1111* type seekRNAs compared to the IS*110* type seekRNA. Though a large 145-170 nt seekRNA was transposase associated, a smaller more abundant seekRNA of 70-90 nt contained the target recognition regions and was shown to be sufficient for transposition in ISEc11. An even shorter region was found to be sufficient for ISEc21 movement. Hence, the precise role of the longer seekRNA remains to be established. In addition, for movement to occur, it was important for the donor IS to be surrounded by the correct target, indicating that the seekRNA is involved in excision of the IS to form the circular intermediate as well as in insertion into the target. Granted the difference observed between the IS*1111* and IS*110* seekRNAs and the known difference relating to the presence of sTIR in the IS*1111* family IS, we conclude that each family was so distinct that it cannot be assumed that the specifics of the movement mechanism are the same in both families.

The IS in the IS*1111* and IS*110* families encode an unusual DEDD transposase type and structure predictions using AlphaFold2 revealed the domain structure for the first time. Though similar overall, there were significant differences in the variable region of IS*1111* and IS*110* ISs and this region appears to interact with the catalytic (DEDD/RuvC) domain in the dimer. How the seekRNA interacts with this protein and which domain binds to the IS ends are not known. Further *in vitro* and structural studies will be needed to resolve these questions.

The importance of systems to reliably introduce large segments of DNA to a specified location in a directional manner has recently been highlighted [31]. The potential for the systems studied here to be exploited for genetic engineering and biotechnology applications is supported by our finding that a cargo flanked by IS ends can be moved and that the target site can be changed by altering both the target-matching stretch in the seekRNA and the sequence flanking the donor. However, the orientation specificity of IS movement that enables orientation-specific insertion will need to be accounted for when selecting a new target. Although future work will be needed to determine the longest and shortest lengths of cargo insert that can be efficiently moved, a clear advantage of the systems reported here over others studied to date [32–35] is the fact that only a single protein of modest size (Tnp are about 340 aa for IS*1111*s and 315-450 for IS*110*s) plus a short seekRNA is required.

## Methods

### Phylogenetic analysis and modelling of IS*110* and IS*1111* family members

The protein sequence of annotated members of the IS*110* and IS*1111* family members (349 sequences) were curated from ISfinder database [1]. Full-length protein sequences that contained the DEDD motif and correct start codon were clustered at 70% identity and one member of each cluster was selected to generate a sequence alignment. Multiple sequence alignment of the selected members of the IS110 and IS111 family members along with Piv comprising a total of 197 sequences was performed on MEGA: Molecular Evolutionary Genetics Analysis version 11 [2] using default settings of Clustal Omega [3]. Phylogenetic tree was generated on Mega using the default settings of neighbour-joining tree and rooted using the midpoint. The multiple sequence alignment (MSA) was further analysed with WebLogo tool to identify conserved residues. A percent identity matrix was created using Clustal Omega. 3D structure prediction of proteins sequences was performed using AlphaFold2 [4] to generate the monomers, dimers and tetramers and visualised on Pymol [5] using ISEc11 and ISEc21 as representative members of the IS*110* and IS*1111* family members. Protein sequence conservation was analysed using Consurf tool [6].

### Genome mining of consensus target site of IS*110* and IS*1111* family members

Sequences for 349 total IS*110* and IS*1111* family members were extracted from ISFinder database [1]. DNA sequences were searched against the preformatted non-redundant nucleotide BLAST database (version date 2022-11-21) using BLAST+ version 2.13.0, restricting to bacterial taxonomic IDs. The BlastN output was filtered by E value of 0, 100% identity, and 95% identity; subject length >=100,000 and <=10,000,000; query coverage 100%. This resulted in the curation of GenBank coordinates where each IS was inserted. From those coordinates and extra 200 base pairs were selected from each side of the identified IS. A custom script was run to query against the new +/-200 and a new database was created of IS+/-200 bp on each side. The flanking sequences were concatenated as pre-insertion sites. All pre-insertion sites for each member of the IS*110* and IS*1111* family members were grouped and filtered after an MSA. Identical sequences were removed in order to generate a non-redundant MSA and unique insertion events. The resulting unique flanking sequences were aligned using Clustal Omega and the MSA was analysed by a custom script to generate a WebLogo [7] to find the consensus target sequence for each IS.

### Molecular cloning and plasmids

*E. coli* strain DH5α was used in this study for molecular cloning and transposition assays. The sequence of selected members of the IS*110* (ISEc21) and IS*1111* (ISEc11, ISKpn4, ISPa11, ISPst6) family members were retrieved from ISfinder, searched using BlastN [8] to locate an insertion point of each member and 100 bps from the flanking sides were added to each IS sequence (IS+100bp) (Supplementary Table 2). Each IS+100bp were synthesised by IDT as a gBlock and the sequences were cloned into pUC19 under the control of lac promoter in BamHI site (Supplementary Table 3). The target sequences were created as described above and the DNA sequences corresponding to the target sites for each IS where ordered as a gBlock from Integrated DNA Technologies (IDT) and cloned into pRSF Duet-1. The Chloramphenicol resistance gene (*cat* gene) containing its endogenous promoter, and the *mCherry* gene were ordered as gBlocks (IDT) and cloned into pCDF Duet-1. All molecular cloning was performed by the Gibson cloning method unless specified otherwise. All plasmid maps and primers are listed (Supplemental Table 3).

### Purification of Tnp RNP complex

The transposase gene was cloned into pCDF-Duet-1 with an N-terminal 6xHis-Maltose-Binding-Protein (MBP), a TEV cleavage site and a C-terminal Strep-TagII. For *in vivo* RNA transcription, the entire IS sequence were cloned into pUC19 under T7 promoter. BL21 (DE3) *E. coli* cells were transformed with either the Tnp plasmid alone or co-transformed with the Tnp plasmid and the corresponding IS plasmids by electroporation using 100 ng of each plasmid. Colonies were grown in 500 mL of LB supplemented with Spectinomycin (100 ug/mL) and Ampicillin (100 ug/mL) until OD_600nm_ of 0.6-0.8 when 0.5 mM IPTG (isopropyl β-D-1-thiogalactopyranoside) was added to the cell culture to induce the expression of the tagged transposase and transcription of the IS and grown for an additional 16 h at 18 °C. The cells were pelleted by centrifugation at 5000 g for 10 min, resuspended in Lysis buffer (100 mM Tris-HCl pH 8.0, 500 mM NaCl, 5% Glycerol, 1 mM TCEP) and lysed by sonication. Cell debris was removed by centrifugation (30 min at 17,000 g), clarified supernatant was loaded onto 1 mL of StrepTactin-XT (high capacity) affinity chromatography column (IBA Lifesciences) and washed with 5 CV of Lysis buffer. The protein was eluted with addition of 50 mM Biotin to the Lysis buffer. The fractions containing the transposase were combined and concentrated using 30 kDa Amicon centrifugal concentrators and stored at -80 °C. The transposase alone and transposases expressed in the presence of the corresponding IS were used for nucleic acid digest gels and small RNA sequencing. Protein samples were prepared as standard analysis for SDS-PAGE by adding NuPAGE™ LDS sample loading buffer and incubated at 98 °C for 10 min to denature, then were loaded on NuPAGE™ 4-12% Bis-Tris Protein Gels (Invitrogen) and electrophoresis was performed at 200 V for 20 mins in MES running buffer (Invitrogen). The gels were stained with Coomassie Brilliant Blue and destained in water. To analyse nucleic acids bound to the transposase, 10 µL of protein alone or co-purified complex with nucleic acids were incubated with either 1 µL of DNase I (18 U/µL) or 1 µL of RNase (10 U/µL) and incubated at 37 °C for 30 min. After the digest, samples were mixed with 2X RNA loading dye (95% formamide, 10 mM EDTA, bromophenol blue, Xylene Cyanol), heated at 98 °C for 2 min to denature before loading on a 7% (19:1) TBE-Urea (7 M) denaturing polyacrylamide gel and electrophoresis was performed at 200 V for 30 mins using 1X TBE running buffer. The gels were stained with SYBR GOLD for nucleic acid visualization.

### Small RNA-Sequencing and analysis

100 µL of the purified transposase alone and Tnp-RNA (RNP) complex was incubated with 5 µL of DNase (20 U/µL) and incubated for 30 min at 37 °C, then 5 µL of Proteinase K was added and incubated for 30 min at 37 °C. Once reaction was complete, the RNA was ethanol precipitated and the pellet washed with 70 % ethanol. The RNA pellet was resuspended in 15 µL MQW. 2 µg of purified RNA was used as total input for preparing RNA libraries for sequencing. Collibri Stranded RNA Library Prep Kit for Illumina Systems (Thermo Fisher Scientific) was used and the libraries were prepared using manufacturer’s protocol and sequenced using an iSeq100 System (Illumina). These reads were then aligned to the sequence of their corresponding IS using the Burrows-Wheeler Aligner alignment tool. The coverage plot was visualised using Integrative Genomics Viewer (IGV) [9] or Tablet v1.21.02.08 [10]. RNA fold prediction was performed with MXfold2 [11].

### Mini-circle formation

Mini-circle formation experiments were conducted using full-length IS sequences+100bp flanking sequences cloned into pUC19. *E. coli* strain DH5α was transformed with plasmid containing the full IS sequence and cultured in LB media for 16 h at 37 °C, then subsequent standard plasmid isolation was carried out (Bioline). DNA template was diluted to 10 ng/µL and 1 µL was used to perform inverse PCR using primers facing outwards from the IS. The corresponding primer set used for each IS are listed in the supplementary material (Supplementary Table 4). PCR was performed in a final reaction volume of 20 µL using Phusion™ reaction buffer, 0.2 mM of dNTPs, 0.5 µM of each primer, and 1 U of Phusion™ polymerase. Thermocycler conditions consisted of denaturation cycle (98 °C, 2 min) followed by 35 cycles of denaturation (98 °C, 15 s), annealing (52 °C, 20 s), extension (72 °C, 20 s), and a final cycle of amplification (72 °C, 40 s). The PCR product was then run on a 1% agarose gel, then isolated using PCR gel extraction kit (Bioline) and Sanger sequenced to identify the minicircle junction product.

### *In vivo* transposition assay

*In vivo* transposition assay was carried out upon co-transforming *E. coli* strain DH5α with the pDonor plasmid containing the entire IS+100bp flanking sequences in pUC19 (Amp resistance) and the pTarget plasmid containing the corresponding target sequence of the IS cloned into pRSF Duet-1 (Kan resistance) (Supplementary Table 3). Co-transformations were performed by electroporation. The transformants were plated onto agar plates and grown overnight at 37 °C. Colonies were grown into 2 mL of LB media supplemented with the antibiotics for 16 h at 37 °C. Standard plasmid isolation was carried out (Bioline). The resulting plasmid DNA templates were diluted to 10 ng/µL and 10 ng was used as template for PCR to detect transposition. Primer pair sets used were complementary to the IS (R) in the *tnp* and the pTarget plasmid backbone (F). The alternate pair (F) on IS and (R) on pTarget plasmid backbone was also performed. The PCR reaction was performed as the standard condition described in the mini-circle formation section, with the following changes, extension (72 °C, 30 s), and final cycle of amplification (72 °C, 60 s). The PCR products were run on 1% agarose gel and the corresponding band was gel extracted using PCR gel extraction kit (Bioline) and further sequenced by Sanger method using one of the PCR primers. The sequencing result was analysed by MAFFT [12] alignments on the target sequence and target sequence+IS left and right ends to confirm that correct insertion had occurred.

### Rescued transposition

A pDonor IS ΔNCR plasmid was generated upon removal of the NCR sequence of the IS cloned into pUC19. A pNCR containing the corresponding NCR sequence of the IS followed by the HDV sequence was cloned under the T7 promoter into pCDF Duet-1 vector. Rescued transposition assay was performed by co-transforming the combination of pTarget and pDonor IS ΔNCR or pTarget, pDonor IS ΔNCR and pNCR (Supplementary Table 3)by electroporation into *E. coli* BL21 (DE3) cells. The transformants were cultured in LB media until OD_600_ of 0.4-0.6 was reached then induced using 0.5 mM IPTG and grown at 25 °C for 16 h before harvesting. Standard plasmid isolation was carried out (Bioline). The resulting plasmid DNA templates were diluted to 10 ng/µL and 10 ng was used as template for PCR to detect transposition. Primer pair sets used were complementary to the IS (R) in the *tnp* and the pTarget plasmid backbone (F). The alternate pair (F) on IS and (R) on pTarget plasmid backbone was also performed. The PCR reaction was performed as the standard condition described in the mini-circle formation section, with the following changes, extension (72 °C, 30 s), and final cycle of amplification (72 °C, 60 s). The PCR products were run on 1% agarose gel and the corresponding band was gel extracted using PCR gel extraction kit (Bioline) and further sequenced by Sanger method using one of the PCR primers. The sequencing result was analysed by MAFFT alignments on the target sequence and target sequence+IS left and right ends to confirm that correct insertion had occurred.

### *In vivo* transposition assay with cargo

*In vivo* transposition assay with cargo was carried out upon co-transforming *E. coli* strain *E. coli* BL21 (DE3) cells with the pDonor plasmid containing the entire IS+100bp flanking sequences in pUC19 (Amp resistance) with the insertion of *cat* at the 5’end of the IS (between LE sequence and the *tnp*, 50 bases after the LE of the IS) or at the 3’of the IS (between the NCR and RE sequence, 46 before the RE of the IS) and the pTarget plasmid containing the corresponding target sequence of the IS cloned into pRSF Duet-1 (Kan resistance). Co-transformations were performed by electroporation. The transformants were cultured in LB media until OD_600_ of 0.4-0.6 was reached then induced using 0.5 mM IPTG and grown at 25 °C for 16 h before harvesting. Standard plasmid isolation was carried out (Bioline) followed by PCR to detect the transposition and mini-circle formation and DNA sequencing to confirm the exact location were performed as described above.

### *In vivo* transposition assay in *trans*

*In vivo* transposition assay in *trans* with cargo was carried out upon co-transforming *E. coli* strain *E. coli* BL21 (DE3) cells with a pDonor, a pTarget and a pSeekRNA plasmids. A pDonor (pUC19) contains the *cat* gene flanked by 100 bp on the 5’ and 100 bp on the 3’end. The 5’ end flanking sequence (L) is composed of the 50 bp on the left side of the insertion point of the IS (LF) in the target site and the first 50 bp of the left end of the IS (LE). The 3’ end flanking sequence (R) is from the last 50 bp of the IS in the right end (RE) and 50 bp of right side of the insertion pint of the IS (RF) in the target site. A pSeekRNA plasmid contains the *tnp* and seekRNA followed by the HDV sequence cloned under T7 promoter 1 and 2 on pCDF Duet-1. Another plasmid combination used had the seekRNA cloned in the pDonor plasmid instead of with the *tnp* in the pCDF duet. Co-transformations were performed by electroporation. The transformants were cultured in LB media until OD_600_ of 0.4-0.6 was reached then induced using 0.5 mM IPTG and grown at 25 °C for 16 h before harvesting. Standard plasmid isolation was carried out (Bioline) followed by PCR to detect the transposition using primers that are complementary to the cargo gene (*cat*) (F primer) and the pTarget plasmid backbone (R primer). The PCR product was sequenced to confirm insertion point.

### mCherry reporter transposition assay in *trans*

A pDonor’ plasmid containing the *tnp* and seekRNA followed by the HDV sequence were cloned under the control of a T7 promoter while a strong T7 terminator sequence was cloned in the 3’of the HDV sequence to prevent downstream gene expression. The mCherry cargo gene is flanked by L and R sequences of the corresponding IS, as described above. The length and sequence of each of these components is listed in the supplementary material (Supplementary Table 3). The pTarget’ plasmid contains the IS target sequence cloned downstream of a T7 promoter. mCherry expression can be detected if transposition has occurred. 100 ng of each plasmid (pDonor and pTarget) were either individually transformed or co-transformed into *E. coli* BL21 (DE3) cells via electroporation. Cells were cultured on agar plates containing Spectinomycin and Kanamycin, incubated overnight at 37 °C in 2 mL LB media. Subsequently, a fresh 2 mL LB medium was inoculated with the initial culture and grown until reaching an OD_600_ of 0.6-0.8, then induced with 0.5 mM IPTG and incubated for 4 hours at 25 °C.

Cell assays involved transferring 100 μL of each culture into a 96-well Corning clear bottom plate. Cell density was quantified via absorbance at 600 nm, while mCherry fluorescence was measured using a TECAN infinite M1000Pro plate reader in bottom reading mode, with excitation at 587 nm and emission at 610 nm, each with a bandwidth of 5 nm, and the optimal gain set to 100%. To adjust fluorescence measurements for cell density, fluorescence per cell (FOD) was calculated as the fluorescence intensity divided by the OD600. The percent increase in mCherry fluorescence attributable to transposition was determined by the formula:

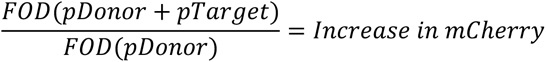

The data was then normalised to the mCherry cargo only samples and represented as percentage increase in mCherry.

This analysis was validated through at least three independent transformations and co-transformations as biological triplicates and sanger sequencing was performed to validate the transposition of the mCherry cargo gene.

### Quantification of mCherry reporter transposition

Transposition frequency was measured by Flow Cytometry using BD LSRFortessa™ X-20 Flow Cytometer. 100 ng of each plasmid (pDonor and pTarget) (Supplementary Table 3) was co-transformed into *E. coli* BL21 (DE3) cells via electroporation. Cells were plated on fresh agar plates containing Spectinomycin, Kanamycin and 0.1 mM IPTG to induce expression of Transposase and seekRNA, as well as expression of mCherry after transposition. The plates were incubated for 16 hours at 37 °C, followed by 4 hours at room temperature. Entire agar plates were scraped containing hundreds of colonies and resuspended and mixed evenly in 1 mL of LB media. Cells were diluted 1 in 2 in PBS (Phosphate Buffered Saline, pH 7.4) and run on the Flow Cytometer. Around 35000-70000 cells were run for each sample until at least 25000 events were recorded, which were gated for single live cells. The cells were counted using forward and side scatter channels, and mCherry fluorescence intensity was detected per cell. The transposition frequency was measured according to the number of single cells with high levels of mCherry fluorescence intensity over total number of single cells. Each set of samples was created by three independent transformations as biological repeats, and the transposition frequency was plotted as bar graphs. Sanger sequencing was performed to validate the transposition of the mCherry cargo gene. Flow cytometry data was analysed using FlowJo™ v10.10 Software (BD Life Sciences) [13].

## Data availability

All data are available in the paper and supplementary information. All code used for genome mining and target sequence consensus creation will be available online at https://github.com/AtaideLab/Targets. Small RNA-Seq data will be available on the NCBI Sequence Read Archive.

## Acknowledgments

We thank the curators of the ISFinder database for continuing to update this valuable resource. This research was supported by the Sydney Informatics Hub and Sydney Analytical, Core Research Facilities of the University of Sydney. We especially thank Cali Willet for the help with the pipeline.

## Author contributions

RMH and SFA conceived the study. SFA, RS, CHP and RMH designed experiments and SFA and RS designed the computational strategy. SFA, RS and RMH performed computational analysis and SFA, RS and CHP performed experiments. SFA and RMH supervised the research. SFA, RS and RMH analysed and interpreted the data. RS and SFA illustrated the manuscript with input from RMH. RMH and SFA wrote the manuscript. All authors reviewed the final manuscript.

## Additional information

**Supplementary information** is available for this paper.

**Correspondence and requests for materials** should be addressed to Dr. Sandro Ataide.

---

**Extended Data Fig. 1.**
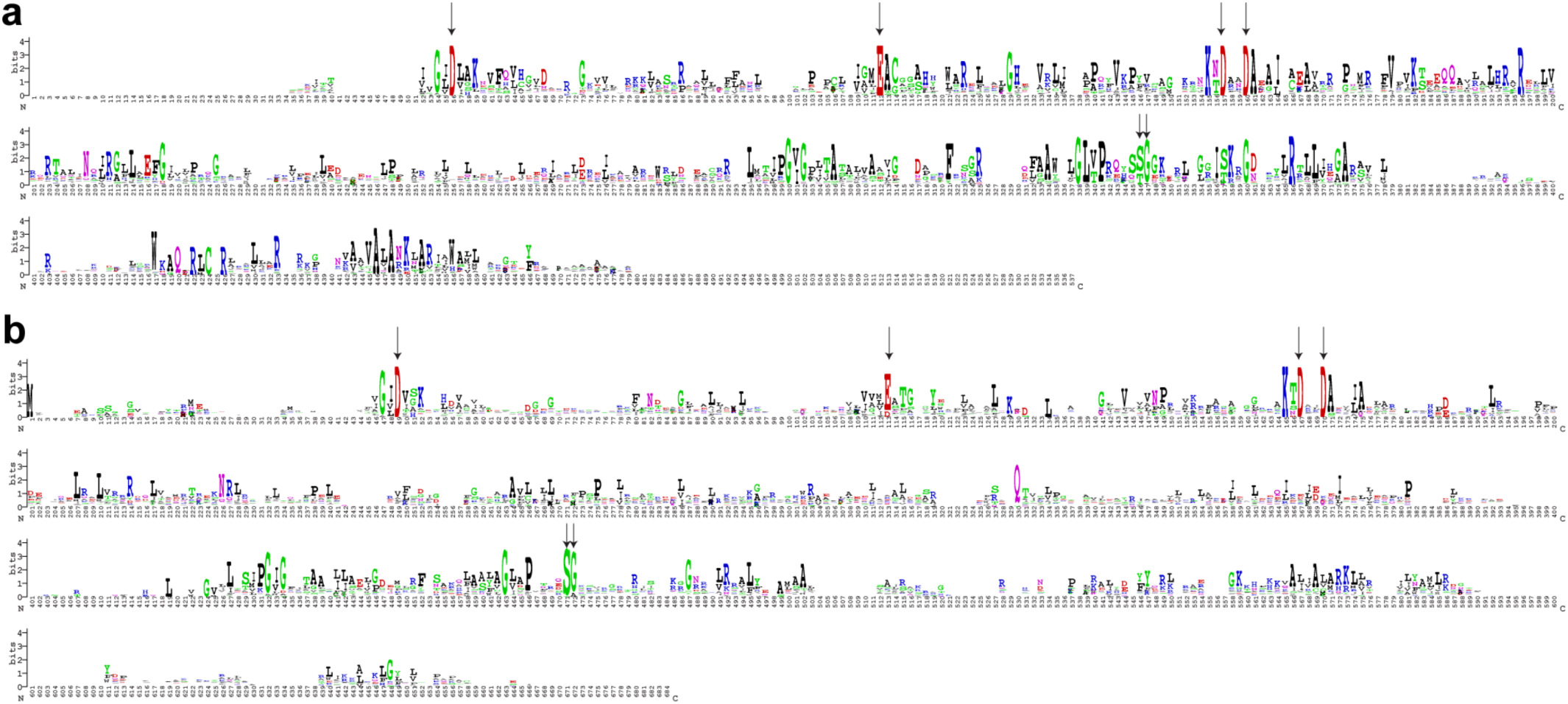
Conserved residues in the transposases from IS*110* and IS*1111* ISs aligned separately. **a.** Transposase sequences from IS*1111* ISs from ISfinder curated and aligned using the Clustal Omega tool in MEGA was used to generate a Weblogo of sequence conservation. Residues are coloured according to their chemical properties: polar (green), basic (blue), acidic (red), hydrophobic (black). Vertical arrows indicate the highly conserved DEDD motif in the N-terminal RuvC domain and SG motif in the C-terminal domain. **b.** Transposases sequences from IS*110* ISs are represented in the same way.

**Extended Data Fig. 2.**
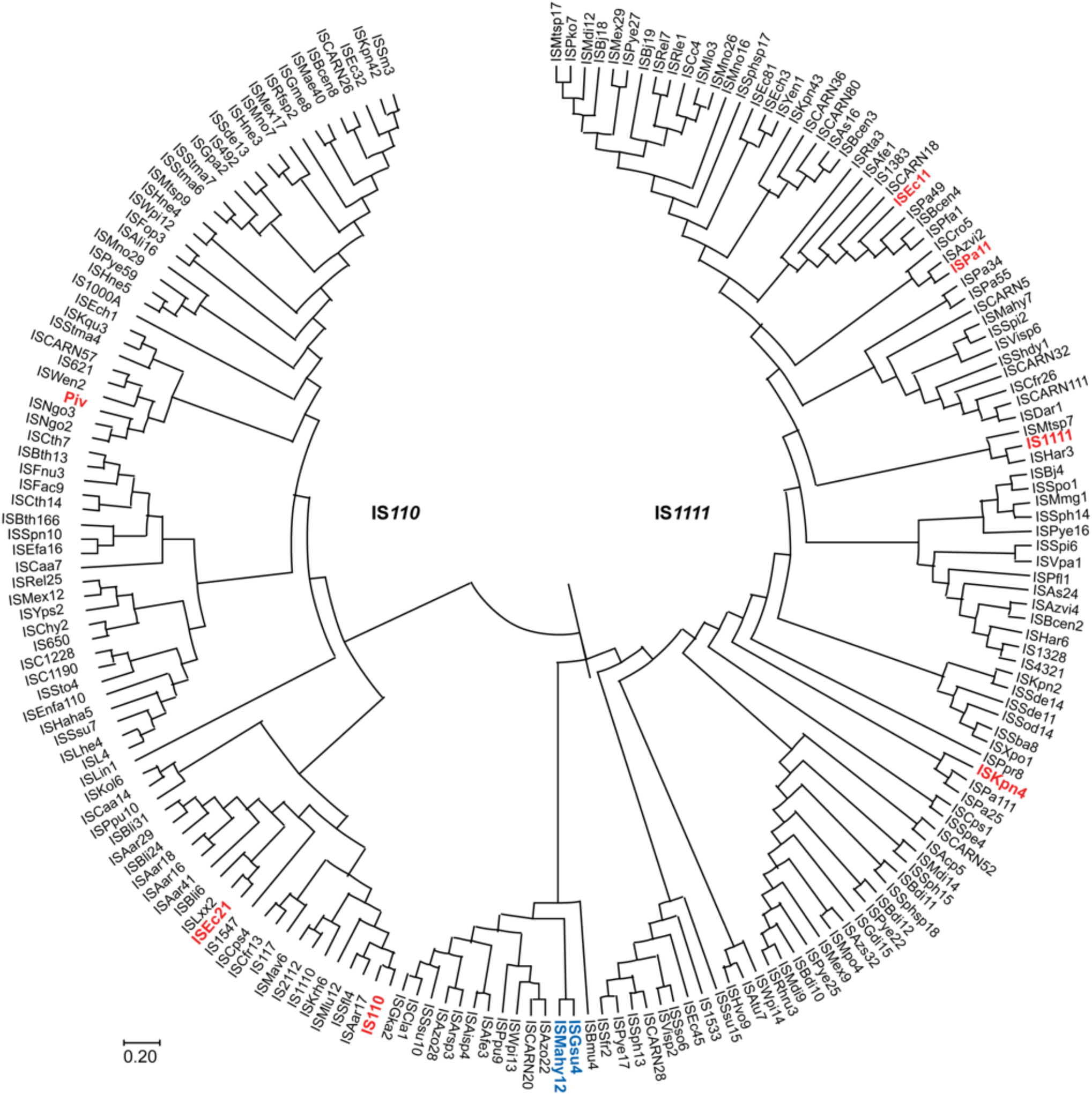
Phylogenetic tree of IS*110* and IS*1111* family transposases. Phylogenetic tree shown as a circle was generated in MEGA using a maximum likelihood neighbour-joining tree with default settings and rooted using the midpoint. The MSA used to generate the tree was comprised of 196 sequences curated from the ISFinder to include a single representative of each cluster of transposases with >70% sequence identity and included the Piv sequence and was aligned using Clustal Omega. Founding family members (IS*110* and IS*1111*), Piv and the ISs used in this study are shown in red. ISs from IS*1111* shown in blue are exceptions that cluster with members of IS*110* family.

**Extended Data Fig. 3.**
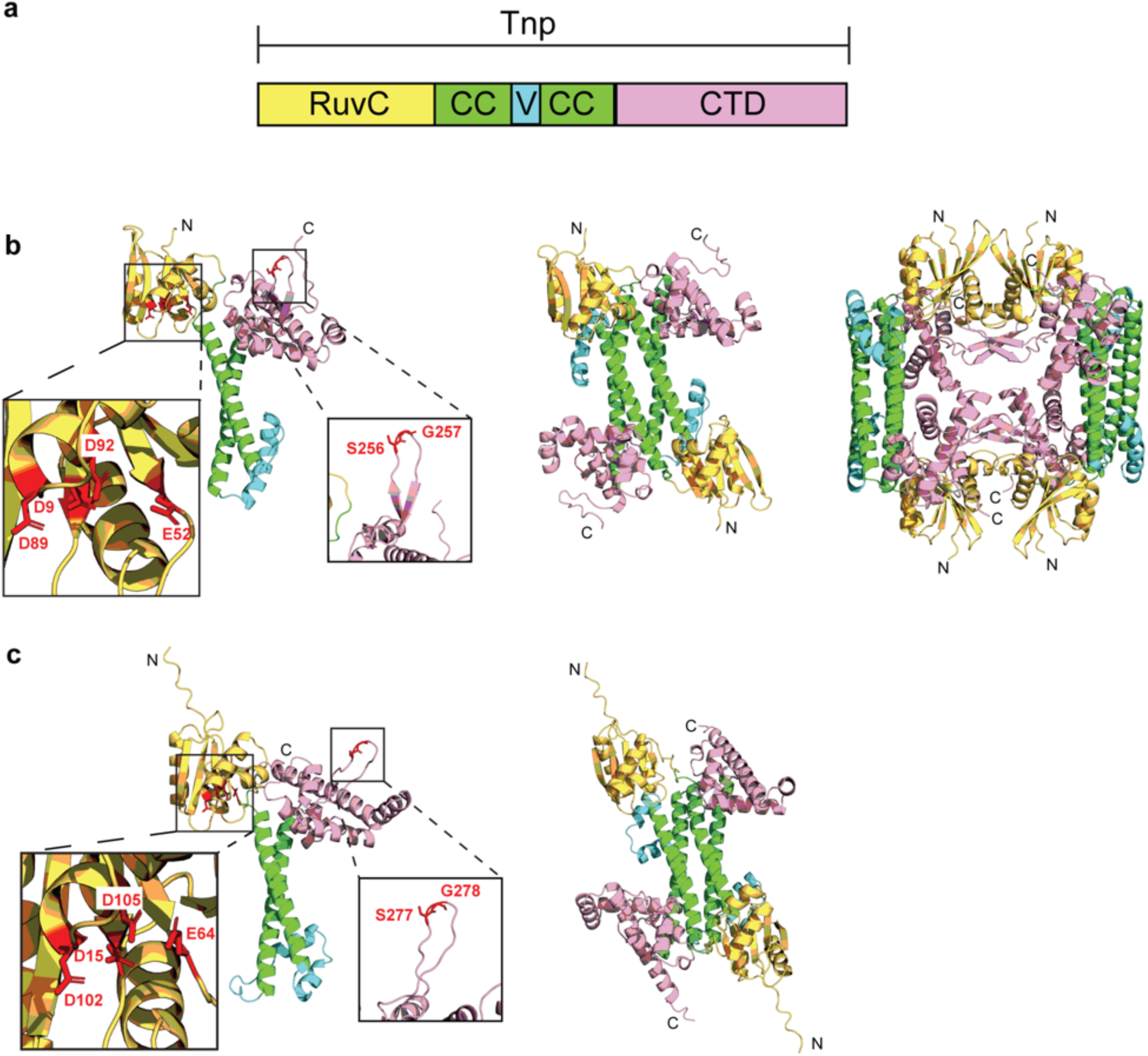
Alphafold2 structure prediction of transposases of IS*1111* and IS*110* IS. **a.** Schematic domain arrangement of Tnp from IS*1111* and IS*110* family members. Domains are colour coded, RuvC in yellow, coiled-coil in green, variable region in cyan and C-term in pink, consistent with the Alfafod2 models in b. **b.** Alphafold2 model of the TnpEc11. Left panel: TnpEc11 monomer with the conserved catalytic amino acids DEDD from the N-terminal domain and SG from the C-terminal domain shown in red sticks in enlargements. Middle panel: Alphafold2 dimer prediction driven by the coiled-coil interactions, placing the variable domain from one monomer next to the RuvC domain of the second monomer. Right panel: Alphafold2 tetramer prediction. **c.** Left panel: Alphafold2 model of the TnpEc21 monomer with the conserved catalytic amino acids DEDD from the N-terminal domain and SG from the C-terminal domain shown in red sticks in enlargements. Right panel: Alphafold2 dimer prediction driven by the coiled-coil interactions, placing the variable domain of a monomer next to the RuvC domain of the second monomer.

**Extended Data Fig. 4.**
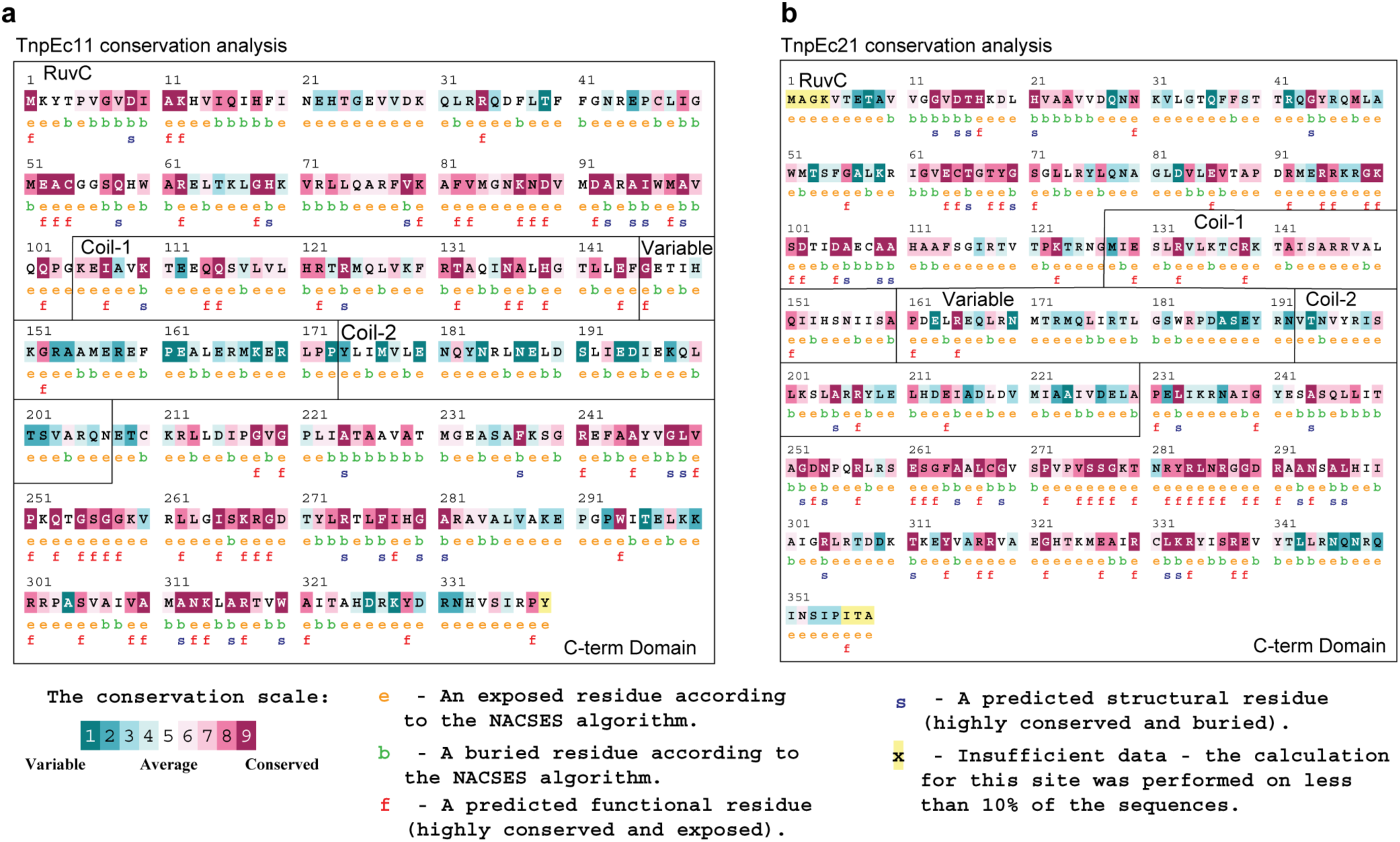
Transposase sequence conservation analysis of ISEc11 and ISEc21. **a.** ConSurf analysis of TnpEc11 as an example of IS*1111* IS with 150 sequences aligned. The transposase domains are indicated with boxes. Keys to conservation scale coding and functional predictions are indicated below. **b.** ConSurf analysis of TnpEc21 as an example of IS*110* IS with 120 sequences aligned. The transposase domains are indicated with boxes. Keys to conservation scale coding and functional predictions are indicated below.

**Extended Data Fig. 5.**
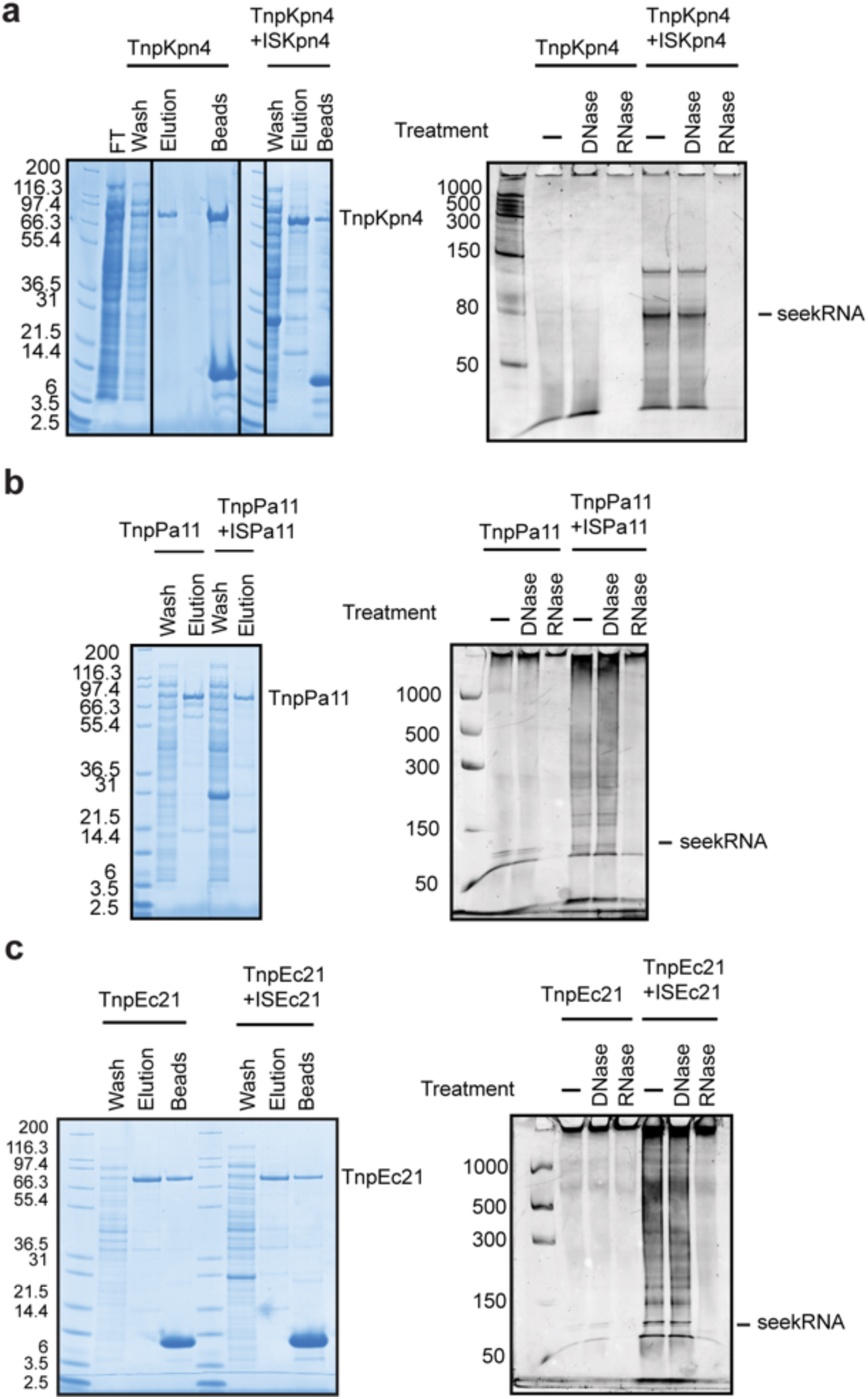
SDS-PAGE and nuclease digest gel analysis. Coomassie stained SDS-PAGE gel shows size and purity of fusion transposases containing an N-terminal 6X-His-MBP tag, and C-terminal StrepTag(II) purified alone or in the presence of their corresponding ISs (see methods and Fig. 2). **a.** TnpKpn4 (85.7 kDa); **b.** TnpPa11 (83.9 kDa); **c.** TnpEc21 (86 kDa) To the right, post-purification, protein samples digested with DNAse or RNAse and resolved in a 7 M Urea polyacrylamide denaturing gel and stained with SYBR GOLD.

**Extended Data Fig. 6.**
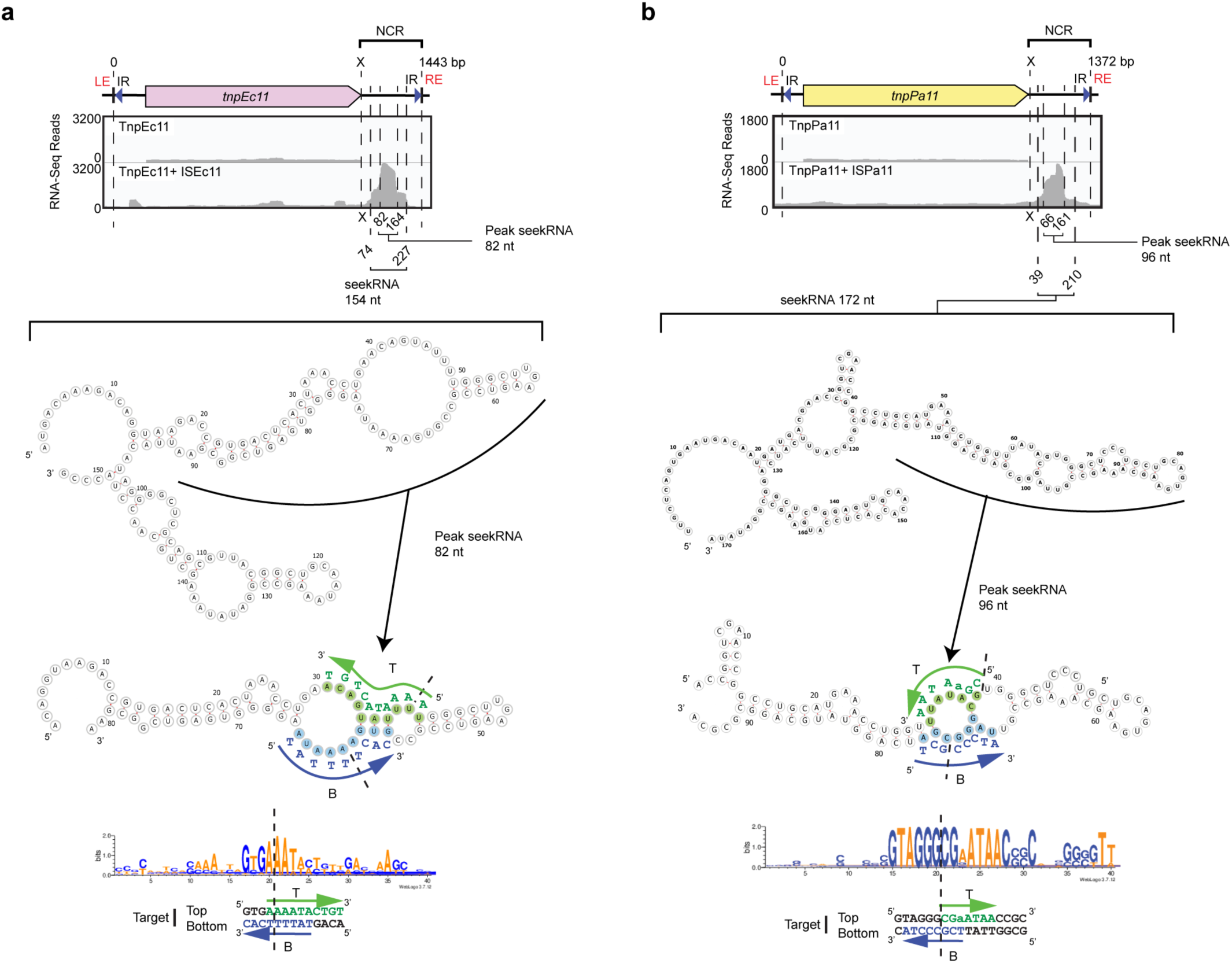
ISEc11 and ISPa11 seekRNA boundaries and folded structure showing target mapping. **a.** Small RNA-seq reads of the TnpEc11 RNP complex aligned to the full length ISEc11 sequence. The long seekRNA (154 nt) spans from 74-227 bp after the *tnp* stop codon (x) and the folded structure prediction shows two hairpin structures with internal loops. The peak or short seekRNA (82 nt) spans from 82-164 bp after the stop codon. On the folded structure bases that are complementary to bases in the top (T) and bottom (B) strands of the target are highlighted in green and blue, respectively with the target sequence alongside. The Weblogo shows the consensus target sequence and below the seekRNA matches are mapped on the target sequence used here, which is complementary mapped onto the peak seekRNA. Arrows indicate the 5’-3’ direction and the insertion point is marked with a vertical dashed line. **b.** Small RNA-seq reads of the TnpPa11 RNP complex aligned to the full length ISPa11 sequence. The long seekRNA (172 nt) spans 39-210 bp after the transposase stop codon (x) in ISPa11. Folded structure prediction shows two hairpin structures with internal loops within each. The peak or short seekRNA (96 nt) spans from 86-161 bp after the stop codon. On the folded structure bases that are complementary to bases in the top (T) and bottom (B) strands of the target are highlighted in green and blue, respectively with the target sequence alongside. The Weblogo shows the consensus target sequence and below the seekRNA matches are mapped on the target sequence used here, which is complementary mapped onto the peak seekRNA. Arrows indicate the 5’-3’ direction and the insertion point is marked with a vertical dashed line.

**Extended Data Fig. 7.**
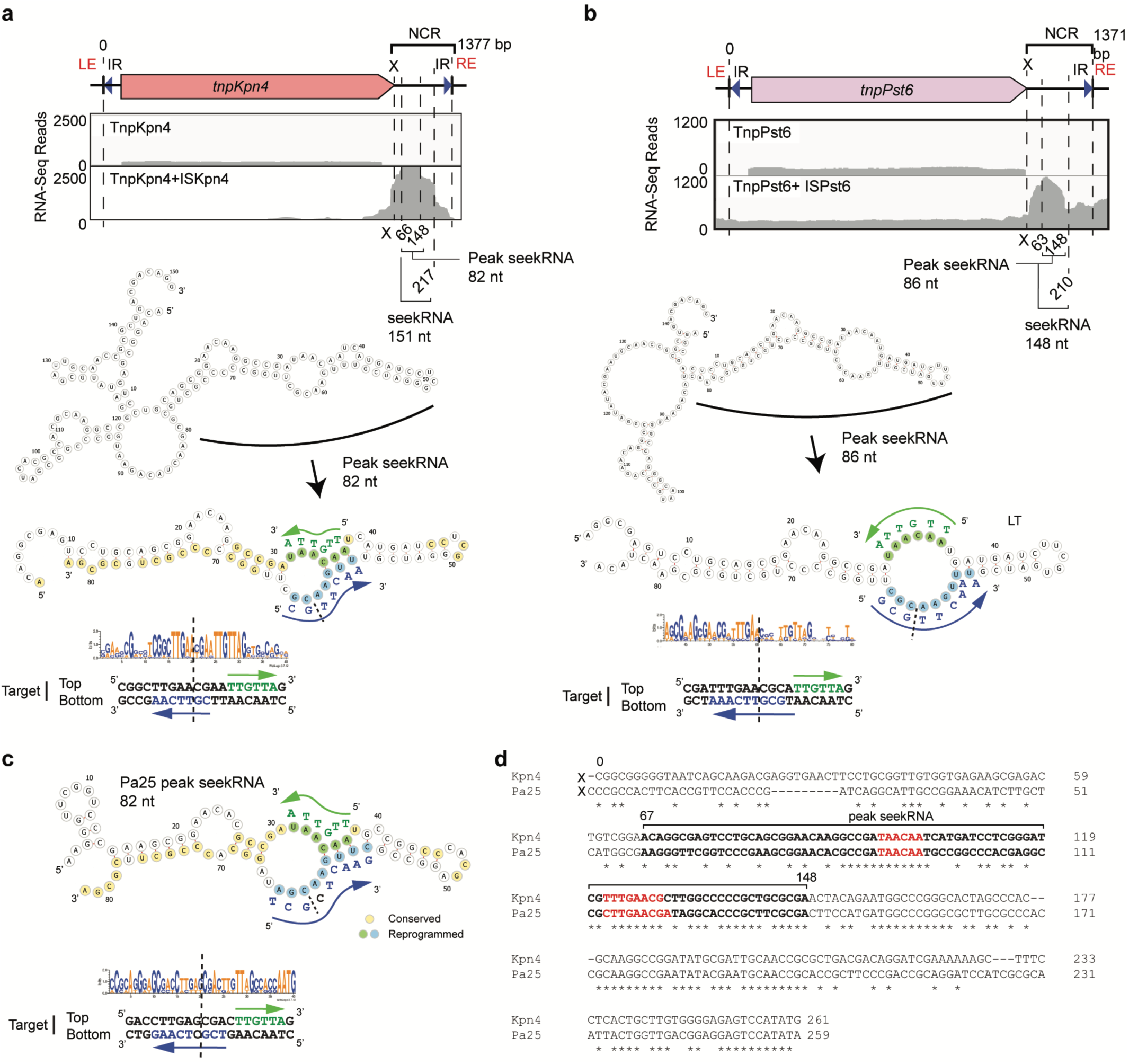
IS*1111* small RNA-seq and seek RNA folded structure similarities and differences compared with their target site. **a.** Small RNA-seq reads of the TnpKpn4 RNP complex aligned to the full length ISKpn4 sequence. The seekRNA is 151 nt in length and peak seekRNA is 82 nt. **b.** Small RNA-seq reads of the TnpPst6 RNP complex aligned to the full length ISPst6 sequence. The seekRNA is 148 nt in length and peak seekRNA is 86 nt. The transposases of ISKpn4 and ISPst6 are 86% identical. **c.** Predicted folded structure of the peak seekRNA of ISPa25 with target site and base pairing sequence to the target top and bottom DNA strands. Bases that are identical in the ISKpn4 seekRNA are highlighted in yellow in both folds. Note: transposases from ISPa25 and ISKpn4 are only 46% identical. **d.** Alignment of the NCR of ISPa25 and ISKpn4 showing identical bases (*) below. The region corresponding to the short seekRNA of ISKpn4 are bold with the target matches in red. In a., b. and c., bases in the folded structure that are complementary to bases in the top (T) and bottom (B) strands of the target are highlighted in green and blue, respectively with the target sequence alongside. The Weblogo shows the consensus target sequence and below the seekRNA matches are mapped on the target sequence. Arrows indicate the 5’-3’ direction and the insertion point is marked with a vertical dashed line.

**Extended Data Fig. 8.**
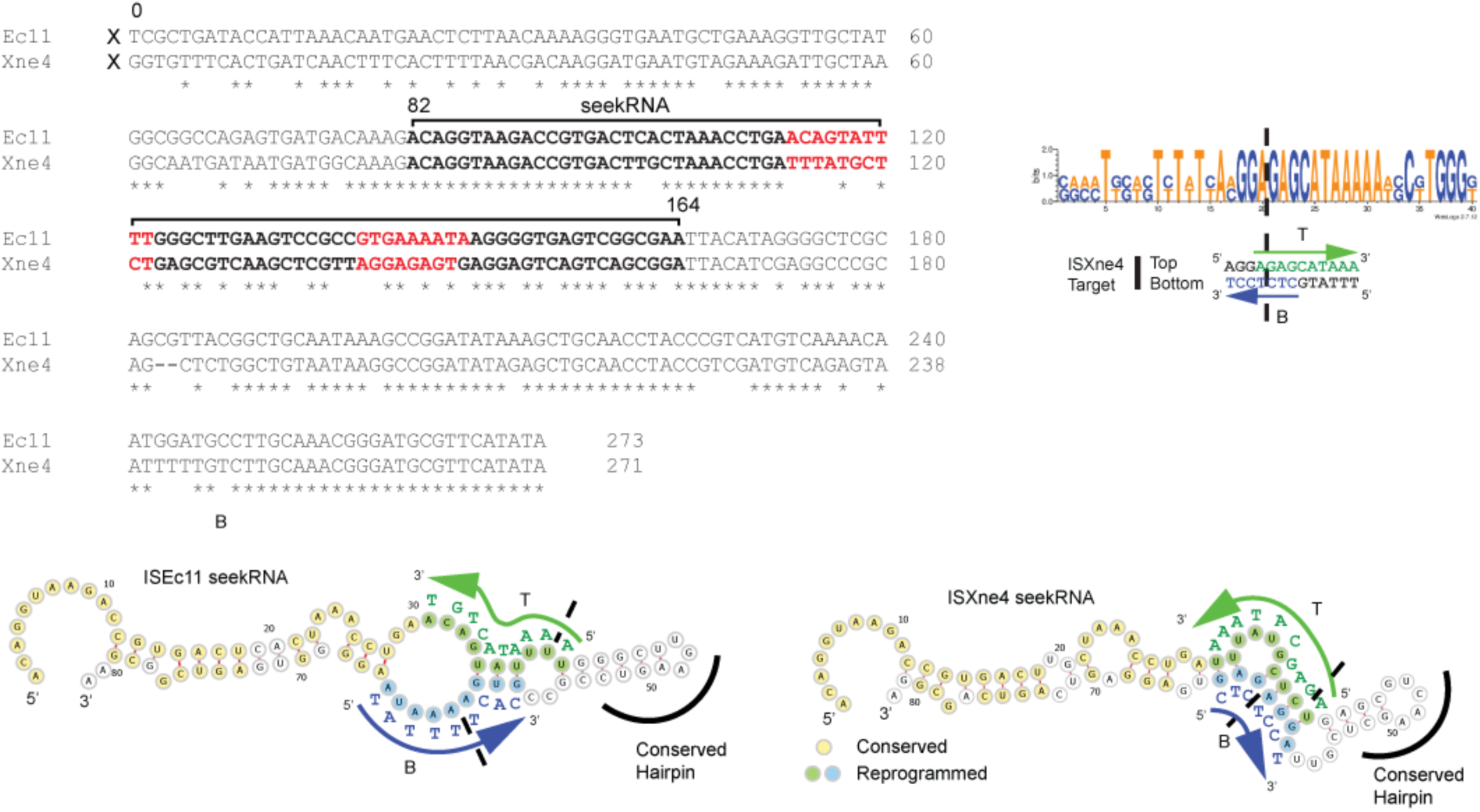
ISEc11 and ISXne4 shows a natural reprogramming. Alignment of the NCR of ISEc11 and ISXne4 starting from the stop codon of *tnp* is shown. Transposases from ISEc11 and ISXne4 are 71.6% sequence identical. The extent of the experimentally determined short (peak) seekRNA for ISEc11 (bp 82-164), and predicted seekRNA for ISXne4 are highlighted in bold with the target matches in red. The target Weblogo showing the consensus target sequence and the target site are shown beside. The similar seekRNA folds are below. Bases in the folded structures that are identical in the ISEc11 and ISXne4 seekRNAs are highlighted in yellow and bases complementary to bases in the top (T) and bottom (B) strands of the target are highlighted in green and blue, respectively. Arrows indicate the 5’-3’ direction and the insertion point is marked with a vertical dashed line.

**Extended Data Fig. 9.**
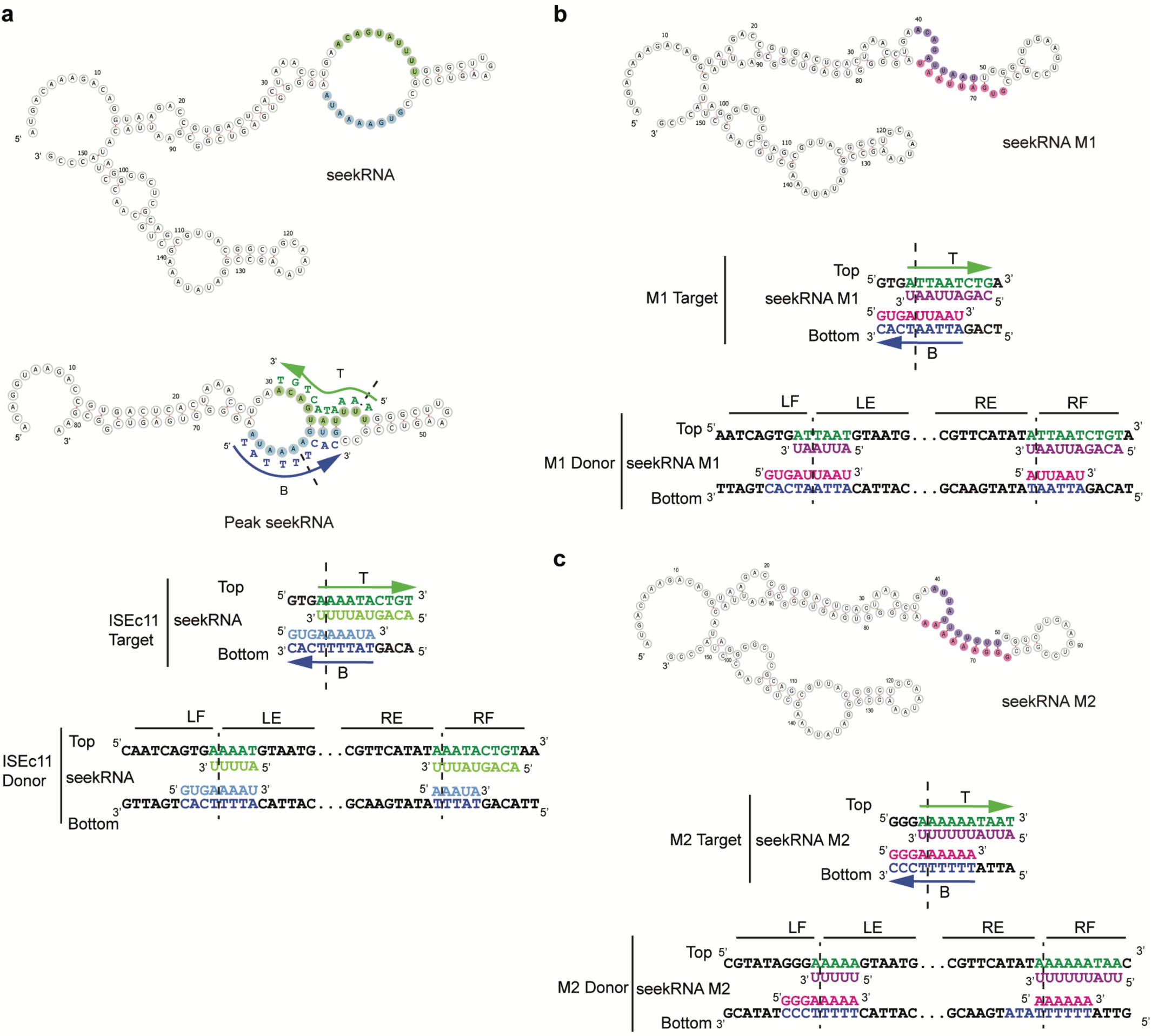
ISEc11 seekRNA modifications for reprogramming into new target site. **a.** Folded structure model of the seekRNA and peak seekRNA of ISEc11 with the bases involved in base pairing with the target site sequence shown in green (top -T) strand and blue (bottom -B) strand. The DNA sequence of the target site is shown for both DNA strands with the arrow pointing towards the 3’ of the corresponding strand and dashed lines indicates the insertion point of the IS. The sequence of the donor LF-LE and RE-RF of ISEc11 can be recognized by the seekRNA with base pairing nucleotides shown in green for top strand and blue for the bottom strand. **b** and **c**. Folded structure model of the reprogrammed seekRNA of ISEc11 with the bases involved in base pairing with the modified target site M1 (b) and M2 (c) showing in purple for the top (T) strand and pink for the bottom (B) strand. The corresponding sequence of the new target site M1 and M22 are shown for both DNA strands with the arrow pointing towards the 3’ of the corresponding DNA strand and dashed lines indicating the insertion point of the IS. The sequence of the donor LF-LE and RE-RF of ISEc11 can be recognized by the seekRNA with base pairing nucleotides shown in green for top strand and blue for the bottom strand.

**Extended Data Fig. 10.**
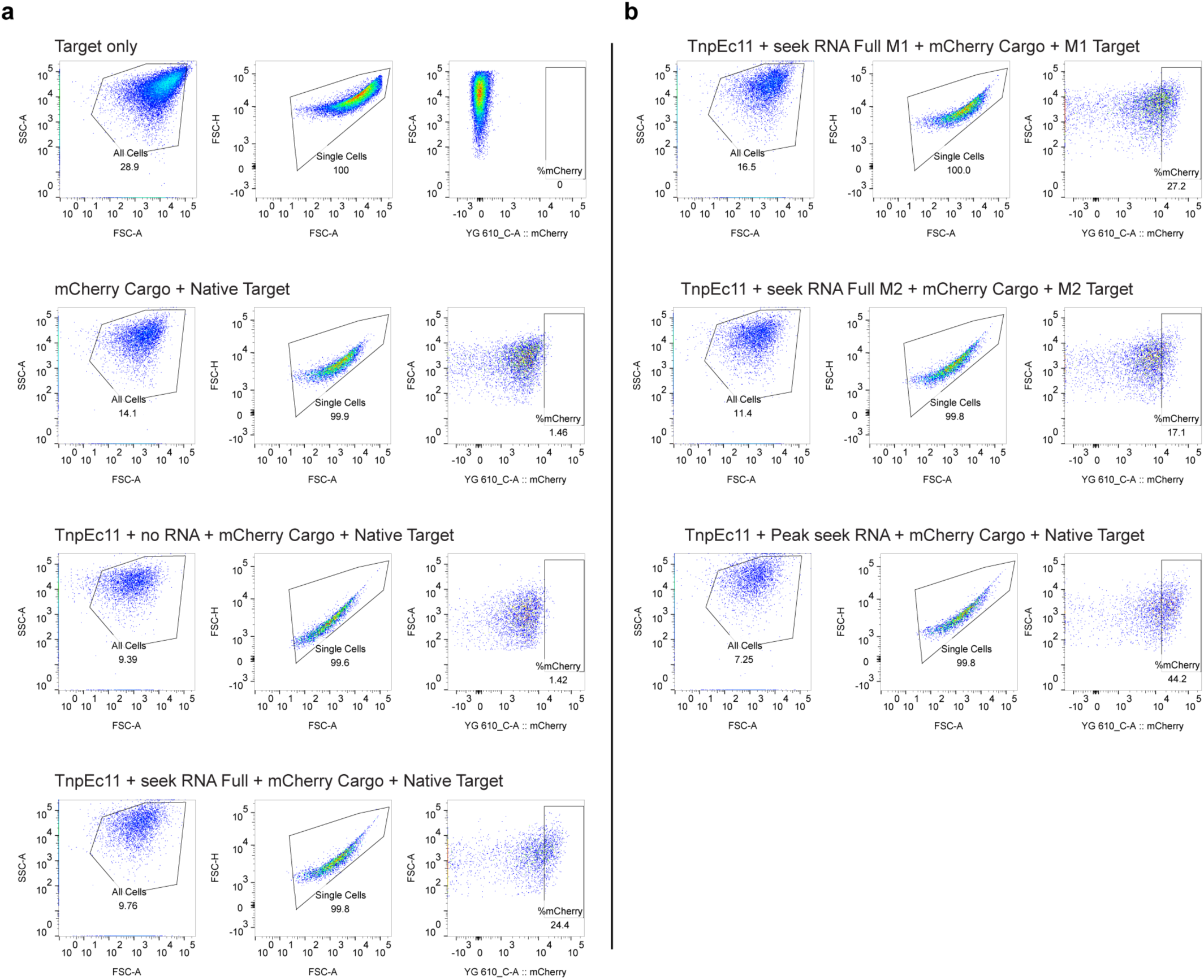
Transposition efficiency of ISEc11 seekRNA and reprogrammed variations using mCherry. **a.** Plots of cells containing a target plasmid (pTarget) and a pDonor and using ISEc11 seekRNA full and mCherry as a cargo (listed above the plots) were gated for all cells (FSC-A and SSC-A), single cells (FSC-A and FSC-H) and mCherry expression shown in each plot. The transposition efficiency of the ISEc11 as the percentage of the total single cells expressing mCherry. **b.** Plots of cells co-transformed with reprogramed pDonor and pTarget plasmids containing a the reprogramed seekRNA and the corresponding target and donor sequence for M1 or M2 and the peak seekRNA with native target of ISEc11gated for all cells, single cells and mCherry expression shown in each plot. The transposition efficiency of the reprogramed ISEc11 and peak seekRNA.

**Extended Data Fig. 11.**
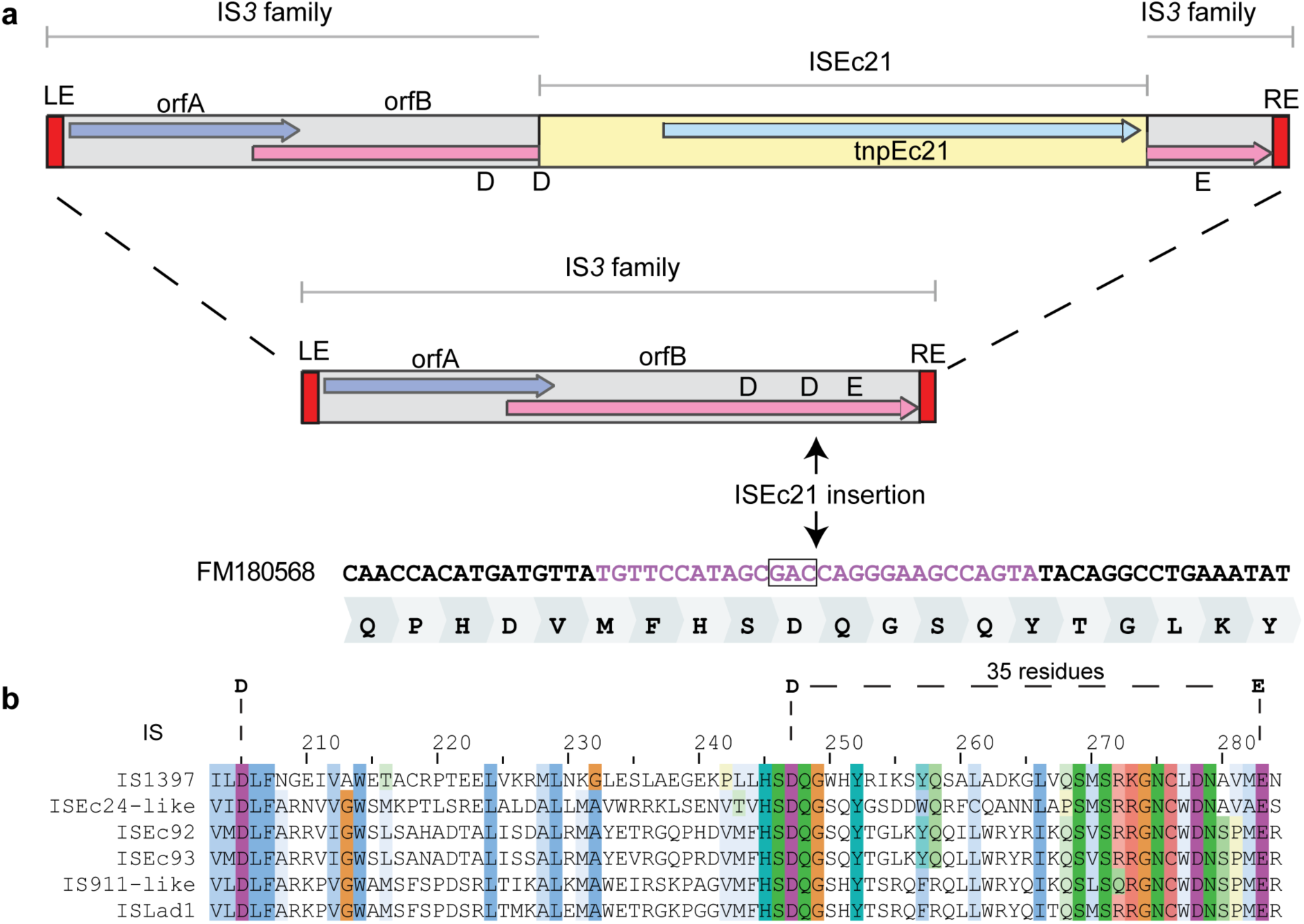
ISEc21 targets the coding sequence of DDE motif of IS*3* members. **a.** Schematic of an IS*3* family IS carrying ISEc21 with the uninterrupted IS below. Coloured arrows indicate the transposase reading frames. The exact location of the ISEc21 insertion in the DNA, after the codon for the conserved D (boxed), is shown with the translation below. A vertical arrow indicates the specific position of the ISEc21. **b.** Multiple sequence alignment of the transposase region of IS where ISEc21 has been found highlighting the aa conservation. Clustal Omega MSA shows clear conservation across multiple residues, including around DDE residues.

**Extended Data Fig. 12.**
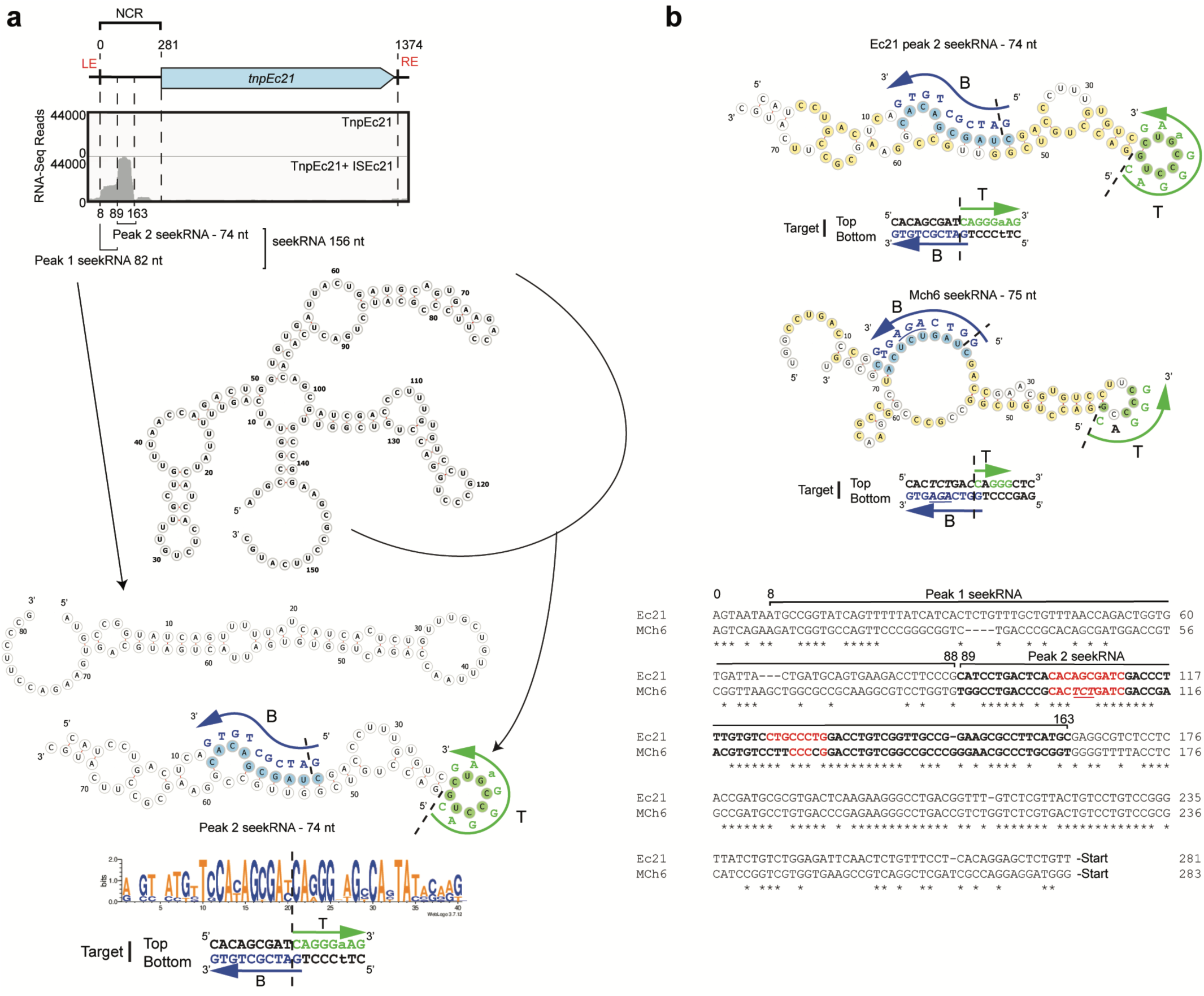
ISEc21 seekRNA folded structures and target site mapping and a naturally occurring reprogramming ISMch6. **a.** Small RNA-seq reads of the TnpEc21 RNP complex aligned to the full length ISEc21 sequence. The full seekRNA folded structure prediction is shown below with folds for the two shorter RNAs below. The peak 1 seekRNA spans from 8-88 bp from the start of the ISEc21. The peak 2 or short seekRNA, is 74 nt long and spans from 89-163 bp. Consensus target sequence mapping shows that the bottom and top strands map to the peak 2 seekRNA. The matches are mapped on the target below as in previous Extended Data Figs. **b.** Comparison of seekRNAs for ISEc21 and ISMch6. Transposases from ISEc21 and ISMch6 are 84% identical. A predicted ISMch6 seekRNA is derived from the alignment of the ISEc21 and ISMch6 NCRs from the start of ISs to the start codon of the transposases. The region of the ISEc21 seekRNA is bold and the target matching regions are red with differences in ISMch6 underlined or black. Other features are as in previous Extended Data Figs.

**Extended Data Fig. 13.**
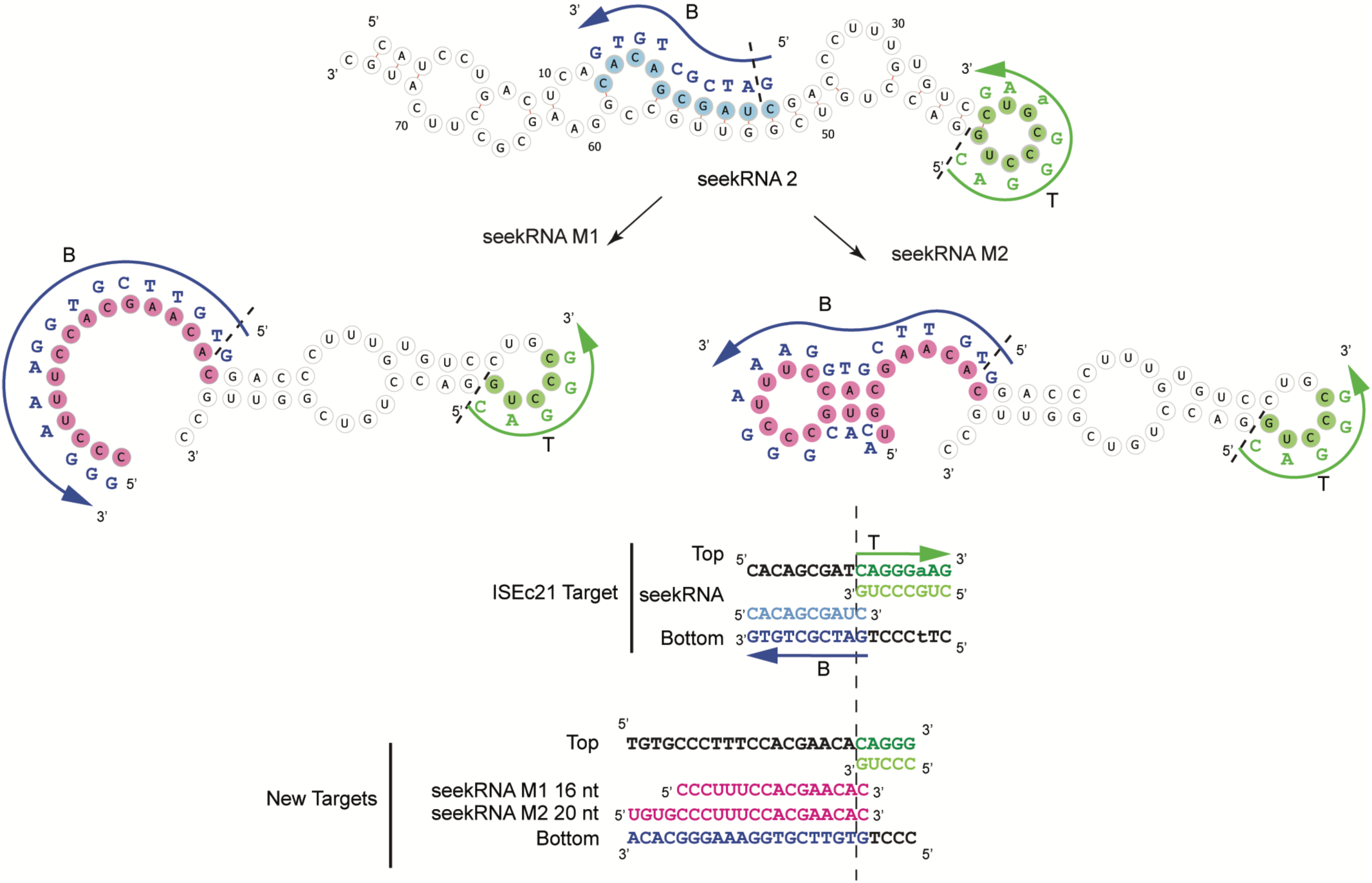
ISEc21 seekRNA modifications for reprogramming to recognize a new target site. The peak 2 seekRNA 74 nt (89-163) of ISEc21 folded structure is shown with the native target site complements shown in blue or green marked on the seekRNA. Reprograming of the seekRNA into M1 and M2 are shown in pink on a shorter 52 nt seekRNA. Addition of 15 nt to the 5’end of the short seekRNA, resulted in a total of 16 nt base pairing as indicated. M2 contains 4 extra nucleotides added to the 5’end of the seekRNA bringing the total base pairing with the target to 21 nt. The new targets are shown below.

